# pHaseMD4AI: Phase-Space Dynamics Dataset with Chemical and pH Perturbations for Physically and Kinetically Consistent Biomolecular AI

**DOI:** 10.64898/2026.07.20.739474

**Authors:** Tiefeng Song, Yixin Guo, Jiahao He, Zheyi Liu, Minying Low, Keying Wang, Yutao Zhang, Zedong Li, Yandong Huang, Yong Wang

**Author notes:** yongwang. These authors contributed equally to this work (Tiefeng Song, Yixin Guo).

## Abstract

Protein function emerges from dynamic conformational ensembles and transitions that are challenging to characterize experimentally and computationally. Recent advances in generative AI have created new opportunities for learning molecular thermodynamics, kinetics, and conformational evolution directly from simulation data, but progress is limited by the availability of large-scale datasets that combine rigorous sampling, complete phase-space information, and diverse physicochemical perturbations. Here, we present pHaseMD4AI, a molecular dynamics dataset that combines a globally equilibrated peptide branch with a protein-scale constant-pH molecular dynamics (CpHMD) branch spanning hundreds of soluble proteins. The peptide branch includes a complete set of canonical tripeptide and tetrapeptide systems together with post-translationally modified (PTM) and protonation-state datasets, providing synchronized atomic coordinates (*R*), velocities (*V*), forces (*F*), and Markov state model-based kinetic annotations. An accompanying web portal (https://isb.zju.edu.cn/md4ai/) enables users to browse, visualize, and download trajectories, annotations, and metadata. As an example application, we demonstrate a sequence-based model that can predict residue-level equilibrium dihedral distributions from sequence. pHaseMD4AI provides a resource for developing and benchmarking molecular machine learning methods while supporting broader studies of biomolecular dynamics under sequence, post-translational modification, and protonation-state perturbations.

## Introduction

Proteins are inherently dynamic systems whose biological functions emerge from thermodynamic ensembles and kinetic transitions rather than single static conformations. ^1,2^ Processes such as folding, molecular recognition, and allosteric regulation are governed by the exploration of complex free-energy landscapes, in which both equilibrium populations and transition pathways play essential roles.^3,4^ Although experimental techniques such as X-ray crystallography and cryo-electron microscopy (cryo-EM) provide high-resolution structural information, they offer only limited access to transient conformational states and molecular motions that underlie biomolecular function. ^1,5^

Molecular dynamics (MD) simulations provide an atomistic framework for investigating biomolecular dynamics by propagating molecular systems according to the underlying physical forces.^6,7^ In principle, sufficiently long simulations can recover equilibrium conformational ensembles together with the corresponding kinetic transition networks. In practice, however, achieving statistically converged sampling remains computationally demanding, particularly for large biomolecular systems where rare conformational transitions occur on long timescales.

To improve sampling efficiency, a wide range of enhanced sampling methods have been developed.^8–11^ These approaches substantially accelerate conformational sampling but often rely on biasing potentials or modified Hamiltonians that perturb the underlying dynamics.^12,13^ Although reweighting and kinetic reconstruction methods can partially recover equilibrium properties,^14–16^ their accuracy depends on the choice of collective variables, bias strength, and sampling completeness.

At the same time, recent advances in machine learning have created new opportunities for modeling biomolecular dynamics. Generative models are increasingly being developed to learn equilibrium conformational distributions, transition pathways, and long-timescale dynamics directly from MD trajectories.^17–27^ The development and benchmarking of these methods require large-scale molecular simulation datasets that combine rigorously sampled equilibrium ensembles with comprehensive physical and chemical annotations.

Protein conformational landscapes are also strongly influenced by changes in chemical state. Post-translational modifications (PTMs), such as phosphorylation, methylation and glycosylation, alter charge distributions, hydrogen-bonding patterns, and steric environments, thereby reshaping free-energy landscapes and regulating protein stability, molecular recognition, and signaling.^28,29^ Likewise, the protonation states of titratable residues vary with the local electrostatic environment and solution pH, coupling directly to conformational transitions, ligand binding, enzymatic catalysis, and membrane transport.^30,31^

Conventional fixed-charge MD simulations assign protonation states before simulation and keep them fixed throughout the trajectory.^32^ Consequently, they cannot explicitly capture protonation-state transitions or their coupling to conformational dynamics under changing pH conditions. CpHMD overcomes this limitation by allowing protonation states to evolve dynamically in response to the molecular environment, enabling simulations of pH-dependent conformational ensembles and proton-coupled molecular mechanisms.^33,34^

As molecular simulation datasets continue to expand, several important limitations remain. Many datasets consist of relatively short simulations or provide only partially sampled conformational ensembles that do not adequately capture metastable transitions.^35–37^ Many widely used datasets primarily distribute atomic coordinates (*R*), while omitting atomic velocities (*V*) and forces (*F*). Because molecular phase space is defined by both coordinates and momenta, and atomic forces encode local gradients of the underlying potential energy surface, the absence of synchronized phase-space information limits downstream development of force-aware and physically consistent machine-learning models.^6,38^ Furthermore, existing datasets rarely integrate sequence variation, PTMs, and protonation-dependent dynamics within a unified resource.

Here, we present pHaseMD4AI, a molecular dynamics dataset that combines globally equilibrated peptide simulations with protein-scale CpHMD simulations. The dataset comprises a peptide branch containing canonical peptides together with PTM and peptide-CpHMD datasets, and a protein-CpHMD branch (Fig. 1). The peptide branch provides synchronized atomic coordinates, velocities, and forces (*R/V/F* s) and Markov state model (MSM)-based kinetic annotations, whereas the protein-CpHMD branch extends the dataset to diverse protonation-dependent folded protein environments through CpHMD simulations. Because achieving global equilibrium is computationally tractable for short peptides but remains prohibitively expensive for large proteins, the two branches provide complementary coverage across different molecular scales. pHaseMD4AI provides a resource for developing and benchmarking machine-learning methods while enabling studies of how sequence variation, PTMs, and protonation-state changes shape biomolecular conformational dynamics.

**Figure 1:**
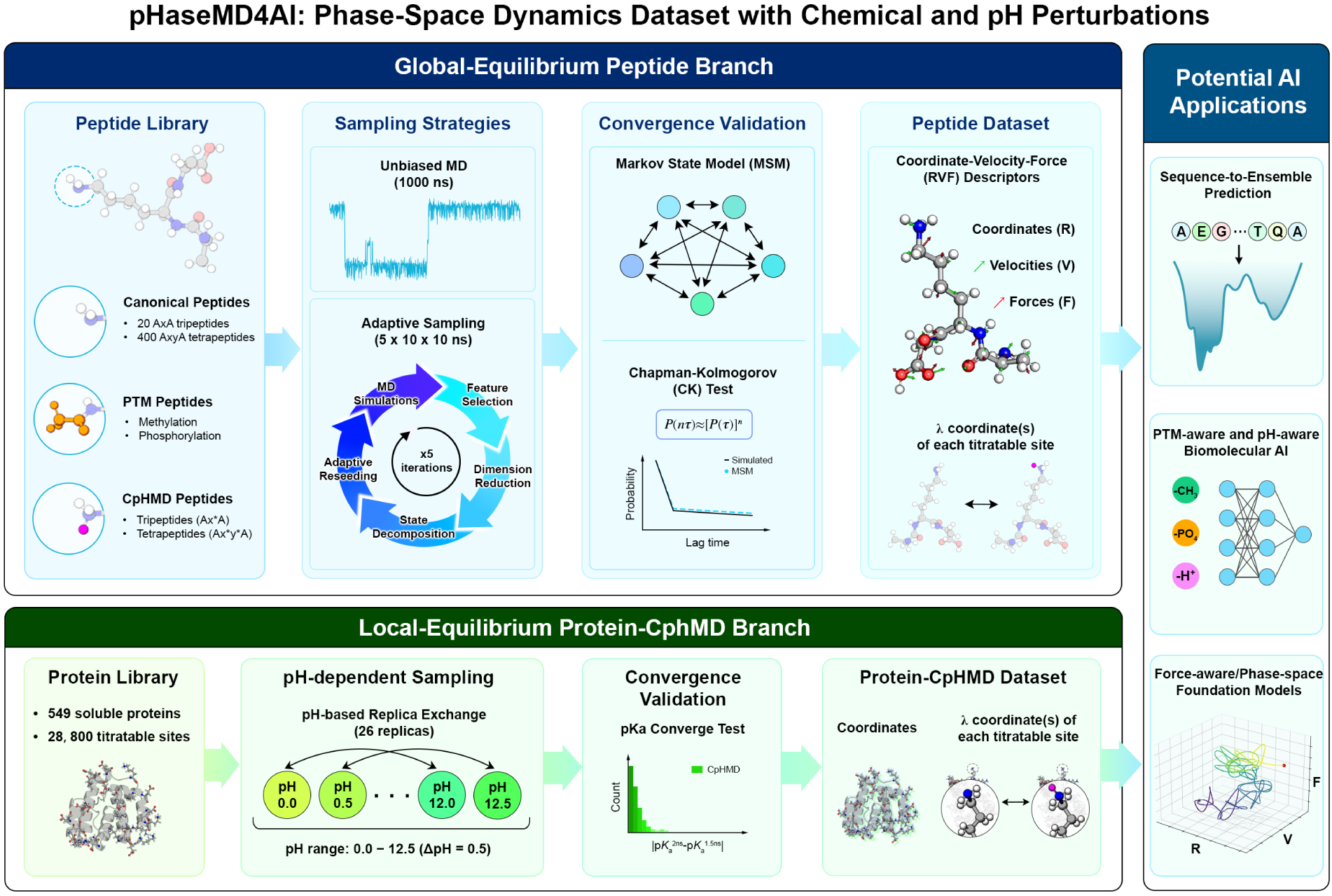
Overview of the pHaseMD4AI dataset architecture and analysis workflow. pHaseMD4AI consists of two complementary branches: a globally equilibrated peptide branch and an exploratory protein-CpHMD branch. The peptide branch comprises a canonical sequence core together with PTM and peptide-CpHMD extensions. Canonical peptides are sampled using long-timescale unbiased MD and adaptive sampling to achieve global equilibrium, PTM systems expand the chemical modification landscape, and peptide-CpHMD simulations capture coupled conformational and protonation-state dynamics. These peptide datasets are further annotated with synchronized *R/V/F* s, MSMs, and Chapman–Kolmogorov (CK) test validation. The protein-CpHMD branch extends the dataset to soluble proteins, providing pH-dependent conformational trajectories and time-resolved protonation-state (*λ*) annotations across diverse folded protein environments.

## Methods and Implementation

### Dataset Architecture

pHaseMD4AI consists of two branches: a peptide branch and a protein-CpHMD branch. The peptide branch contains canonical peptides together with PTM and peptide-CpHMD datasets, whereas the protein-CpHMD branch contains CpHMD trajectories for soluble proteins.

#### Canonical Peptide Dataset

The canonical peptide dataset consists of 20 tripeptides (*H*-Ala-*x*-Ala-OH) and 400 tetrapeptides (*H*-Ala-*x*-*y*-Ala-OH), where *x* and *y* represent the 20 canonical amino acids. These peptides serve as the reference dataset for comparisons with PTM and peptide-CpHMD systems.

All peptides were parameterized using the CHARMM36m force field^39^ and solvated in explicit TIP3P water.^40,41^ Counterions were added to neutralize the systems. After energy minimization and equilibration, production simulations were performed in the NVT ensemble at 300 K. Hydrogen mass repartitioning (HMR) technique ^42^ was applied to enable a 4 fs integration time step. The simulation protocol and validation of the HMR scheme are described in the Supporting Information (Fig. S1).

#### PTM Peptide Dataset

The PTM peptide dataset was constructed from selected tripeptides to investigate the effects of common PTMs on peptide conformational dynamics. The dataset includes methylated Lys and Arg together with phosphorylated Ser, Thr, and Tyr. Lys was modeled in eight methylation states, whereas Arg was represented by two commonly observed methylation forms. Each phosphorylated peptide was simulated in two protonation states (Fig. 2). Initial structures were generated using CHARMM-GUI and subsequently processed using the same simulation workflow and force field as the canonical peptide dataset to ensure consistency across all peptide systems.^43^

**Figure 2:**
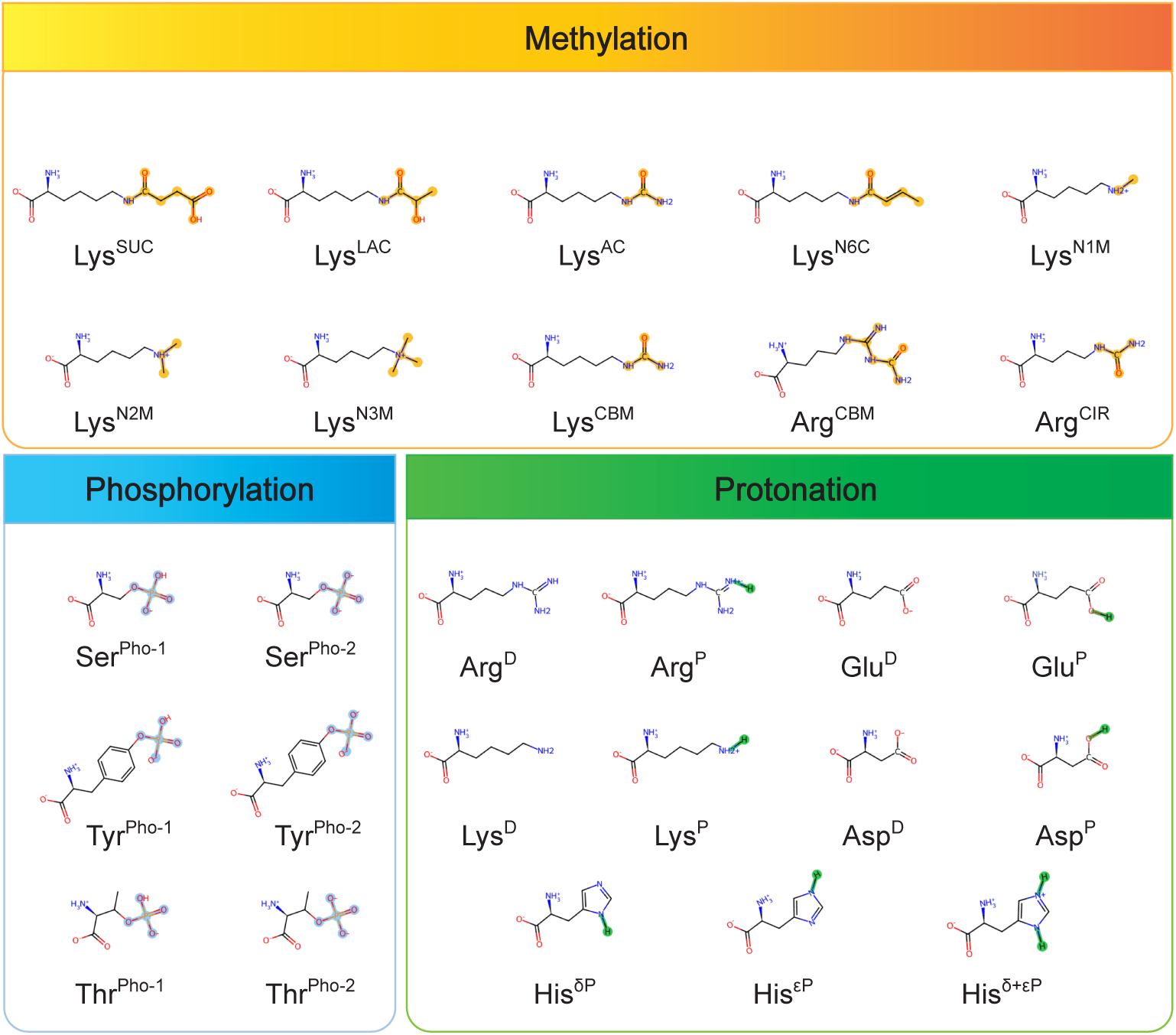
Methylation, phosphorylation, and protonation entries and 2D structural diagrams involved in pHaseMD4AI.

#### Peptide-CpHMD Dataset

To represent protonation-state-dependent conformational dynamics explicitly, a peptide-CpHMD branch was constructed for titratable peptides. Five tripeptides containing titratable residues (Asp, Glu, His, Lys, or Arg) and 25 tetrapeptides representing all pairwise combinations of these residues were included (Fig. 2). CpHMD simulations were carried out at pH 7.0 using a modified implementation of GROMACS 2021 with the *λ*-dynamics approach. ^33^ Protonation-state parameters and topology modifications were generated using pHbuilder.^44^ In this CpHMD method, *λ* is a continuous titration coordinate that describes the protonation state of a titratable residue, where *λ* = 0 corresponds to the fully protonated state and *λ* = 1 corresponds to the fully deprotonated state. The *λ* value evolves continuously between 0 and 1, governed by the Hamiltonian equation. Under constant-pH conditions, protonation states evolve together with the atomic coordinates, so the resulting trajectories capture both structural rearrangements and protonation-state transitions within the same biomolecular system. Unless otherwise specified, the remaining simulation parameters were kept as consistent as possible with the standard pHaseMD4AI workflow. For each titratable site, the dataset records the *λ* coordinate(s) at every saved frame, enabling analysis of time-resolved protonation-state dynamics alongside conformational motion.

#### Protein-CpHMD Branch

To extend pHaseMD4AI from global-equilibrium peptide systems to protein-scale pH-dependent behavior, we constructed a protein-CpHMD branch spanning 549 soluble proteins.^45^ These proteins are sequence-non-homologous, water-soluble proteins randomly sampled from the TR6614 dataset, and for multimers we only simulated their first subunit.^46^ To accelerate simulations, this branch was generated using the GPU-accelerated continuous CpHMD implementation in Amber with the ff14SB protein force field and the GBNeck2 generalized-Born implicit-solvent model (‘igb=8’), together with ‘mbondi3’ intrinsic Born radii and an ionic strength of 0.15 M. Asp, Glu, His, and Lys were treated as titratable residues, whereas Arg was not included in the dynamic titration set. Each system was equilibrated at pH 7.0 and 300 K, and production sampling was performed using pH-based replica exchange over pH 0.0–12.5 at 0.5-pH-unit intervals, corresponding to 26 replicas per protein.^47,48^ Bonds involving hydrogen atoms were constrained with SHAKE, allowing a 2 fs integration time step, and replica-exchange attempts were made every 1000 MD steps. Following the baseline protocol, each replica was typically propagated for 2 ns. To assess short-timescale local equilibrium under this high-throughput regime, we computed the absolute difference between the p*K_a_* estimates obtained at 1.5 and 2.0 ns and used the criterion |Δp*K_a_*| *<* 0.2 to define local equilibrium.

#### Stored Data

For each peptide system, pHaseMD4AI stores synchronized atomic *R*, *V*, and *F* together with kinetic information derived from MSMs. For peptide-CpHMD simulations, the dataset additionally includes the time-resolved *λ* coordinate(s) of each titratable site. For the protein-CpHMD dataset, the released data consist of atomic coordinate trajectories together with the corresponding *λ* coordinate(s) for all titratable sites.

### Global-Equilibrium Sampling of Peptide Systems

Globally equilibrated peptide ensembles were generated using a combination of 1 *µ*s long-timescale unbiased MD simulations and 500 ns adaptive sampling. This workflow was applied to the canonical peptide, PTM peptide, and peptide-CpHMD datasets. Sampling convergence was subsequently assessed using the CK test.

#### Adaptive Sampling

Adaptive sampling was performed to improve sampling of low-population conformational states and rare conformational transitions. The workflow consisted of the following steps:

1. **Initial Sampling.** Each system was initialized from a representative starting conformation and subjected to 10 parallel unbiased MD simulations of 10 ns each.
2. **Feature Representation and Dimensionality Reduction.** Pairwise distances between backbone heavy atoms were used to describe conformational variability. The resulting features were projected onto a low-dimensional space using time-lagged independent component analysis (TICA),^49,50^ which captures the slow collective motions governing long-timescale dynamics.
3. **State Decomposition.** The reduced TICA space was partitioned into microstates using k-means clustering.^51^ MSMs were then constructed from the discretized trajectories, and metastable macrostates were identified using the Perron cluster-cluster analysis (PCCA+) algorithm.^52^
4. **Adaptive Reseeding.** To promote exploration of under-sampled regions, new starting conformations were selected from low-population macrostates. Ten such conformations were used to initiate the next round of simulations.
5. **Iterative Exploration.** After initial sampling, the cycle of feature representation, TICA projection, MSM construction, PCCA+ analysis, and adaptive reseeding was repeated for four iterations. Each adaptive sampling campaign generated approximately 500 ns of aggregate simulation data (10 trajectories × 10 ns × 5 iterations), substantially expanding sampling space with much more affordable cost.

### Kinetic Modeling and Equilibrium Validation for Peptide Datasets

To enable consistent analysis of peptide simulations generated using different sampling protocols, all peptide trajectories were processed using the same MSM workflow.

#### Markov State Model Construction

Peptide trajectories generated by unbiased MD and adaptive sampling were featurized using pairwise distances between backbone heavy atoms. TICA was then performed with a lag time of 100 ps, and the first 10 time-lagged independent components (tICs) were retained for subsequent analysis.

The reduced conformational space was discretized into microstates using k-means clustering. The number of clusters was determined according to the empirical scaling relationship, which accounts for the number and length of trajectories.^53^ MSMs were constructed using a lag time of 100 ps to estimate the transition probability matrix *P* (*τ*), where *P_ij_*(*τ*) denotes the probability of transitioning from state *i* to state *j* over the lag time *τ*. Microstates were further grouped into metastable macrostates using the PCCA+ algorithm. ^52^ By default, each system was partitioned into 10 macrostates; if this decomposition was not feasible, five macrostates were used instead.

#### Convergence Validation

The Markovian behavior of each peptide system was evaluated using the CK test,^51^ which compares transition probabilities predicted by the MSM with those observed directly from the simulations. Only peptide systems that passed the CK test were included in the globally equilibrated peptide dataset. This criterion was applied to the canonical peptide, PTM, and peptide-CpHMD datasets.

The protein-CpHMD dataset was not subjected to CK test validation because the simulations were designed to sample pH-dependent conformational changes in folded proteins rather than to construct globally converged kinetic models.

#### Thermodynamic and Kinetic Characterization based on MSM

We calculated the thermodynamic and kinetic characterizations based on MSMs constructed using unbiased MD and adaptive sampling, respectively. The FES was projected onto the first two tIC dimensions using Koopman-reweighting based on the MSM. ^54^ Mean first-passage times (MFPTs) and reactive fluxes between PCCA+ defined metastable macrostates were then computed to quantify the kinetics of conformational transitions.^51,55–57^

### Representative Analyses

#### Comparison of Ramachandran Distributions Across Peptide Systems

To compare backbone conformational preferences across peptide systems, Ramachandran distributions were analyzed for canonical peptides together with their PTM and peptide-CpHMD counterparts. The tripeptide dataset contained 41 systems (20 canonical peptides, 16 PTM peptides, and five peptide-CpHMD systems), whereas the tetrapeptide dataset contained 425 systems (400 canonical peptides and 25 peptide-CpHMD systems).

For tripeptides, pairwise similarity was quantified using the Jensen–Shannon (JS) divergence between the Ramachandran distributions of the central residue. For tetrapeptides, similarity was calculated from the joint Φ-angle distribution of the two internal residues, which captures the dominant backbone coupling within the tetrapeptide.

Hierarchical clustering was performed on the resulting pairwise JS-distance matrices using average-linkage agglomerative clustering with optimal leaf ordering.

#### Prediction of Dihedral Angle Distributions from Sequence

As an example application of the dataset, we trained a model to predict residue-resolved dihedral angle distributions from peptide sequence. The model was trained using the globally equilibrated canonical peptide dataset, including the non-terminal residues of 20 tripeptides and 400 tetrapeptides. PTM and CpHMD peptides were not included because the current implementation does not explicitly represent modified residues or protonation states.

For each residue, the model predicts both the backbone Ramachandran distribution and the side-chain torsional distributions.

For the evaluation presented here, the 20 tripeptides were used for training, whereas the 400 tetrapeptides served as an independent test set. The model consisted of two stages. First, a variational autoencoder (VAE)^58^ employing circular one- and two-dimensional convolutional layers learned a latent representation of the periodic dihedral distributions. Second, frozen embeddings from a protein language model,^59^ together with residue identity and physicochemical descriptors, were used to predict the latent variables from sequence. The predicted latent variables were then decoded into residue-resolved dihedral distributions using the fixed VAE decoder (Fig. S2).

Prediction accuracy was evaluated on the tetrapeptide test set using reconstruction crossentropy and JS similarity^60^ between the predicted and reference distributions. A detailed description of the model architecture, input features, and training hyperparameters is provided in the Supporting Information.

### Web Platform Implementation

The frontend was developed with Next.js (v14.2.5) using TypeScript (v5.5.4) and integrates the NGL WebGL^61^ molecular viewer (v2.4.0) to render molecular structures and trajectory frames in the browser. The backend exposes RESTful endpoints using FastAPI^62^ (Python v3.10.19) and was served via Uvicorn, while NGINX was used as a reverse proxy to route requests between the web application, API endpoints, and static assets. Trajectory streaming was enabled through MDsrv^63^ (v0.3.5), allowing progressive retrieval of coordinate frames without downloading full trajectories upfront.

## Results

### Dataset Overview

pHaseMD4AI consists of two complementary branches (Fig. 1). The peptide branch contains canonical peptides together with PTM and peptide-CpHMD datasets, providing globally equilibrated conformational ensembles for short peptides. The protein-CpHMD branch extends the dataset to pH-dependent conformational sampling in folded protein environments.^45^

The peptide branch includes synchronized *R*, *V* and *F*, FES, MSM macrostate populations, MFPTs, reactive fluxes, representative structures of metastable states, and equilibrium validation. Whereas the protein-CpHMD branch provides protein trajectories together with time-resolved *λ* coordinate(s) for all titratable residues.

### Global-Equilibrium Peptide Branch Dataset and Representative Analyses

#### Sequence, PTM, and Protonation-State Diversity

The peptide branch comprises three complementary components that capture sequence variation, PTMs, and protonation-state dynamics. The canonical peptide dataset provides comprehensive coverage of tripeptide and tetrapeptide combinations and serves as the reference for subsequent analyses. The PTM extension introduces methylation and phosphorylation variants, enabling direct comparison with the corresponding unmodified peptides. The peptide-CpHMD extension further incorporates protonation-state fluctuations through time-resolved *λ* coordinates recorded alongside conformational trajectories.

For all globally equilibrated peptide systems, pHaseMD4AI provides synchronized R/V/Fs, together with MSM-based kinetic annotations. These data provide a unified representation for systematic analysis of sequence-, PTM-, and protonation-dependent conformational landscapes.

#### Comparison of Ramachandran Landscapes Across Peptide Systems

Ramachandran distributions were compared across canonical peptides, PTM peptides, and peptide-CpHMD systems using hierarchical clustering based on JS divergence (Fig. S3). The clustering and FESs show that PTM and peptide-CpHMD systems generally remain close to their corresponding canonical peptides, indicating that most perturbations alter conformational preferences without producing entirely distinct backbone conformations (Fig. S3-S9).

Among the tripeptides, Gly and Pro formed two well-separated clusters. Excluding Gly and Pro, peptides containing Trp, Val, and Ile formed a distinct cluster, whereas Asp together with protonated Asp (D*) and phosphoserine clustered separately from the remaining amino acids.

PTM variants of Lys and Arg clustered near their corresponding canonical peptides, indicating relatively modest changes in backbone conformational preferences following methylation. Similarly, most titratable residues remained close to their corresponding canonical residues, with His being the main exception.

Phosphoserine showed the largest deviation among the PTM systems and clustered closer to Asp and protonated Asp than to unmodified serine, consistent with the introduction of a negatively charged phosphate group.

A similar trend was observed for tetrapeptides (Fig. S10). Gly- and Pro-containing sequences again formed distinct clusters, whereas the remaining peptides exhibited a more complex organization owing to coupling between adjacent residues. Peptide-CpHMD systems remained close to their corresponding canonical sequences, indicating that protonation primarily reshapes the populations of existing conformational states.

#### Prediction of Dihedral Angle Distributions from Sequence

To demonstrate one potential application of the dataset, we trained a sequence-based model to predict residue-resolved equilibrium dihedral angle distributions using the canonical peptide dataset. On the independent tetrapeptide test set, the predicted backbone Ramachandran distributions and side-chain *χ*_1_–*χ*_4_ distributions closely matched those obtained from MD simulations and approached the reconstruction accuracy of the latent-space autoencoder (Fig. S11-S12), indicating that equilibrium residue-level conformational preferences can be inferred directly from peptide sequence.

### Protein-CpHMD Branch Dataset

The protein-CpHMD branch contains 549 soluble proteins comprising 28,800 titratable sites (Fig. S13).^45^ These simulations extend the dataset from short peptides to folded protein systems spanning a wide range of structural environments.

The dataset was generated using short CpHMD simulations designed to sample protonation-dependent conformational changes across many proteins. For each protein, the released data include coordinate trajectories together with the time-resolved *λ* coordinate of every titratable site.

### Comparison with Existing Molecular Dynamics Datasets

We compared pHaseMD4AI with representative molecular dynamics datasets, including long-timescale simulation datasets, large-scale protein repositories, and specialized databases (Table 1). Existing resources differ substantially in their scope and design. Early datasets, such as the D. E. Shaw trajectories,^64,65^ provide exceptionally long unbiased simulations for a limited number of fast-folding proteins, whereas repositories such as MoDEL,^66^ Dynameomics,^67^ ATLAS,^68^ mdCATH,^69^ and DynaRepo^70^ emphasize broader structural coverage across diverse biomolecular systems. Other resources, including MemProtMD, GPCRmd, SCoV2-MD, and ManyPeptidesMD, focus on specific classes of biomolecules, such as membrane proteins, GPCRs, viral proteins, or peptides.^71–74^

pHaseMD4AI complements existing molecular dynamics resources by combining globally equilibrated peptide simulations with a protein-scale CpHMD dataset within a unified framework. The peptide branch provides synchronized atomic R/V/Fs, and MSM-based kinetic annotations for canonical peptides together with datasets covering PTMs and protonation-state dynamics. In particular, the PTM datasets include methylation and phosphorylation variants that enable systematic comparisons between modified and unmodified peptides, whereas the peptide-CpHMD datasets capture the coupling between conformational dynamics and protonation-state changes. The protein-CpHMD branch further extends the resource to 549 soluble proteins simulated using pH-replica exchange, providing protonation-state sampling across diverse folded protein environments. Together, the two branches integrate sequence variation, PTMs, and protonation-state dynamics across peptide- and protein-scale molecular systems.

**Table 1:**
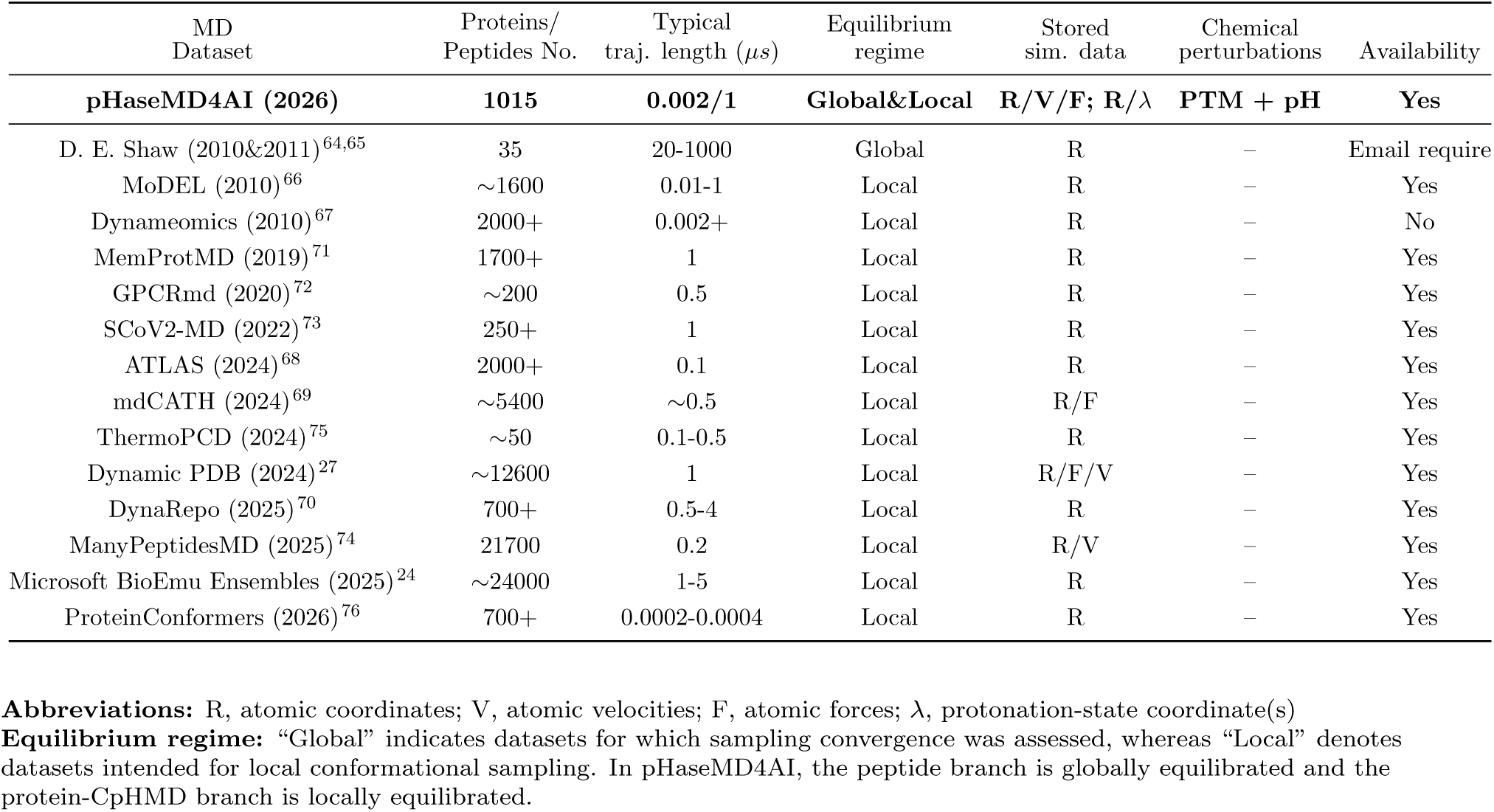
Comparison of pHaseMD4AI with Existing Molecular Dynamics Datasets.

| MD Dataset | Proteins/<br>Peptides No. | Typical<br>traj. length ($\mu s$) | Equilibrium<br>regime | Stored<br>sim. data | Chemical<br>perturbations | Availability |
| --- | --- | --- | --- | --- | --- | --- |
| **pHaseMD4AI (2026)** | **1015** | **0.002/1** | **Global&Local** | **R/V/F; R/$\lambda$** | **PTM + pH** | **Yes** |
| D. E. Shaw (2010&2011)<sup>64,65</sup> | 35 | 20-1000 | Global | R | – | Email require |
| MoDEL (2010)<sup>66</sup> | ~1600 | 0.01-1 | Local | R | – | Yes |
| Dynameomics (2010)<sup>67</sup> | 2000+ | 0.002+ | Local | R | – | No |
| MemProtMD (2019)<sup>71</sup> | 1700+ | 1 | Local | R | – | Yes |
| GPCRmd (2020)<sup>72</sup> | ~200 | 0.5 | Local | R | – | Yes |
| SCoV2-MD (2022)<sup>73</sup> | 250+ | 1 | Local | R | – | Yes |
| ATLAS (2024)<sup>68</sup> | 2000+ | 0.1 | Local | R | – | Yes |
| mdCATH (2024)<sup>69</sup> | ~5400 | ~0.5 | Local | R/F | – | Yes |
| ThermoPCD (2024)<sup>75</sup> | ~50 | 0.1-0.5 | Local | R | – | Yes |
| Dynamic PDB (2024)<sup>27</sup> | ~12600 | 1 | Local | R/F/V | – | Yes |
| DynaRepo (2025)<sup>70</sup> | 700+ | 0.5-4 | Local | R | – | Yes |
| ManyPeptidesMD (2025)<sup>74</sup> | 21700 | 0.2 | Local | R/V | – | Yes |
| Microsoft BioEmu Ensembles (2025)<sup>24</sup> | ~24000 | 1-5 | Local | R | – | Yes |
| ProteinConformers (2026)<sup>76</sup> | 700+ | 0.0002-0.0004 | Local | R | – | Yes |

**Abbreviations:** R, atomic coordinates; V, atomic velocities; F, atomic forces; $\lambda$ , protonation-state coordinate(s) **Equilibrium regime:** “Global” indicates datasets for which sampling convergence was assessed, whereas “Local” denotes datasets intended for local conformational sampling. In pHaseMD4AI, the peptide branch is globally equilibrated and the protein-CpHMD branch is locally equilibrated.

### An Interactive Web Platform

To enhance the accessibility, analytical capability, and practical utility of pHaseMD4AI, we developed a comprehensive web-based platform (https://isb.zju.edu.cn/md4ai) that provides integrated data exploration, visualization, and analysis services for MD datasets (Fig. 3, S14-S21). The platform enables researchers to interactively inspect conformational ensembles, free energy landscapes, kinetic transitions, and phase-space descriptors without requiring local data preprocessing or specialized MD software. The frontend further synchronizes molecular structures with time-series properties, such as potential energy, through shared frame and time indices, enabling users to examine the structural context of energetic fluctuations and dynamically navigate between trajectory and property views.

**Figure 3:**
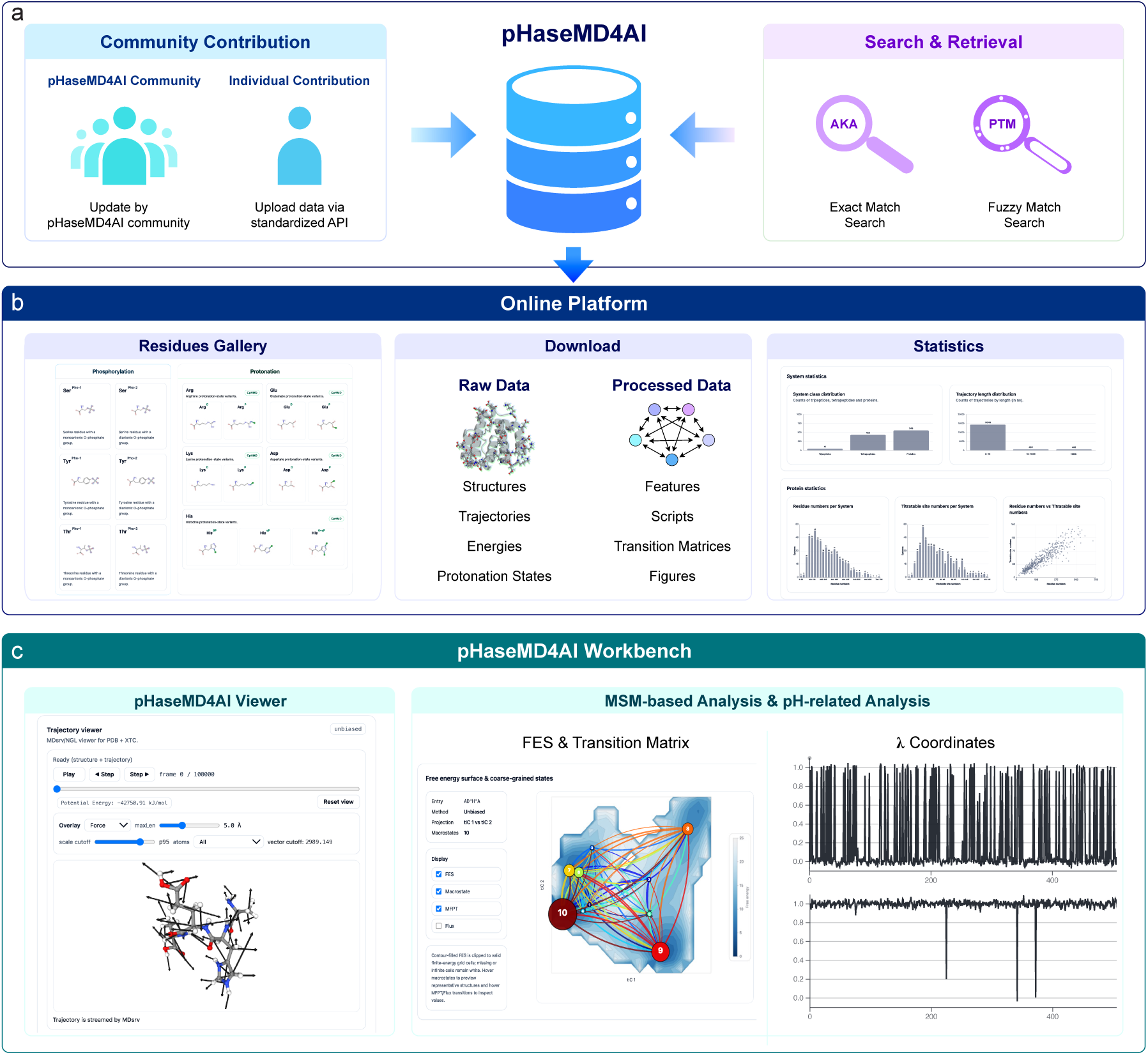
pHaseMD4AI online platform framework. (a) pHaseMD4AI is an online resource platform for storing, streaming, and analyzing MD simulation data of short peptides and water-soluble proteins, incorporating both user contributions and centralized updates from the developers. The platform features intuitive search algorithms for comprehensive database filtering as well as fuzzy search capabilities that allow users to search for specific sequence types. (b) Users can search for systems based on visualized residue types, download multi-level information and gain an overview of the entire database. (c) The workbench supports interactive visualization (pHaseMD4AI Viewer) for simulations data, alongside analyses based on MSM and protonation states.

The platform also functions as the integration layer for both the global-equilibrium peptide branch and the exploratory protein-CpHMD branch. Users can browse canonical peptide entries, PTM variants, peptide-CpHMD systems, and protein entries through the same interface while retaining access to branch-aware metadata, equilibrium diagnostics for peptides, and branch-specific metadata for proteins. This unified presentation is important because it exposes pH-dependent effects across scales rather than presenting the different dataset branches as isolated resources. It also facilitates comparisons among canonical sequences, chemically modified variants, and alternative protonation states, thereby helping users identify how specific physicochemical perturbations reshape conformational behavior. The search layer correspondingly supports exact retrieval by sequence or protein name together with modification-aware fuzzy retrieval, enabling direct navigation between canonical entries and their chemically perturbed variants. In addition, the platform provides graphical representations of canonical residues, their corresponding chemical modifications, and the alternative protonation states, with residue-type-based retrieval supporting systematic exploration of chemically and protonationally related systems. These features collectively lower the technical barrier to accessing and interpreting the dataset and provide an interactive entry point for both molecular simulation studies and AI model development.

#### Kinetic and Thermodynamic Analysis Tools

The platform provides a deep dive into the analysis based on trajectories and MSMs. Key features include:

- **Trajectory Data:** Each molecular trajectory is accompanied by time-resolved potential energy and *λ* coordinate(s) (where applicable), enabling integrated analysis of conformation, energetic, and protonation-state dynamics across peptides and proteins.
- **Macrostate Distribution:** Interactive network graphs display the equilibrium populations of kinetically distinct states identified via PCCA+ and visualize the transfer rates (*k_ij_*) between macrostates.
- **Representative Conformations:** For peptide macrostates, users can inspect representative structural ensembles and compare their branch-specific contexts, including PTM state or protonation annotations.
- **Equilibrium Diagnostics:** The platform hosts CK test results for global-equilibrium peptide systems so that users can examine consistency between predicted and observed transition probabilities before downstream reuse.

#### Multi-tiered Data Download Interface

Recognizing the diverse needs of the computational biology community, pHaseMD4AI offers a multi-tiered data access model:

1. **Trajectory Level:** Processed peptide files containing coordinates, forces, velocities, and perturbation-aware trajectories, as well as protein-CpHMD coordinates with time-resolved *λ* coordinate(s).
2. **Representative Structure Level:** Users can retrieve representative macrostate structures in PDB format for peptide systems together with branch-specific protonation annotations.
3. **Model Level:** TICA projections, transition times and fluxes, macrostates populations and equilibrium diagnostics are available for researchers focusing on kinetic modeling and pH-sensitive analysis.

By centralizing these tools, the pHaseMD4AI website serves as both a data repository and a diagnostic suite, facilitating AI model development and biological hypothesis generation across global-equilibrium peptide dynamics and local-equilibrium protein pH landscapes.

## Discussion

pHaseMD4AI adopts a complementary two-branch design. The peptide branch emphasizes globally equilibrated conformational ensembles with synchronized phase-space information, including atomic coordinates, velocities, and forces together with equilibrium validation and kinetic annotations. In contrast, the protein-CpHMD branch prioritizes broad coverage of pH-dependent conformational behavior across hundreds of folded proteins. This strategy combines rigorous equilibrium characterization where computationally feasible with largescale exploration of protonation-dependent protein dynamics.

A distinguishing feature of pHaseMD4AI is its explicit representation of molecular dynamics in phase space. Unlike most existing MD datasets that primarily provide coordinate trajectories, pHaseMD4AI synchronizes coordinates, velocities, and forces throughout globally equilibrated peptide simulations. Together with MSM-derived kinetic models and equilibrium validation, these data provide simultaneous structural, dynamical, and energetic information, supporting applications ranging from force matching and neural force fields to physics-informed generative models of molecular dynamics. By incorporating PTMs and protonation-state dynamics within the same framework, the dataset also enables systematic investigation of how chemical perturbations reshape free-energy landscapes, redistribute conformational ensembles, and alter molecular kinetics.

The two branches serve complementary scientific purposes. Within the peptide branch, canonical peptides provide a globally equilibrated reference dataset, while PTM and peptide-CpHMD simulations expand the chemical and protonation-state diversity of conformational ensembles. Cross-system comparisons show that both PTMs and protonation-state changes redistribute equilibrium populations within a shared statistical mechanical framework. In contrast, the protein-CpHMD branch is intended to capture diverse protonation-dependent conformational behaviors across soluble proteins rather than construct globally converged kinetic models. These simulations therefore provide locally equilibrated ensembles suitable for studying pH-dependent conformational changes, residue protonation behavior, and diverse electrostatic environments.

The sequence-based prediction of residue-resolved dihedral distributions demonstrates one proof-of-concept application of the dataset. Using only peptide sequence information, the model accurately recovered equilibrium backbone and side-chain dihedral distributions for previously unseen tetrapeptides, illustrating that globally equilibrated phase-space data can support learning equilibrium conformational ensembles directly from sequence. The inclusion of PTM and CpHMD simulations further provides opportunities to develop future models that explicitly account for chemical modifications and protonation-state changes.

Beyond the examples presented here, pHaseMD4AI provides a general resource for data-driven studies of biomolecular dynamics. The synchronized phase-space information, equilibrium conformational ensembles, kinetic annotations, and perturbation-aware simulations establish a foundation for developing physics-informed machine learning models that jointly learn molecular structure, dynamics, kinetics, and thermodynamics. Potential applications include benchmarking generative molecular dynamics methods, learning free-energy landscapes and conformational kinetics, evaluating adaptive sampling strategies, and investigating PTM-dependent and pH-dependent structural dynamics.

Despite its broad coverage, pHaseMD4AI has several limitations. Global equilibrium was established only for the peptide branch, whereas the protein-CpHMD simulations prioritize biological diversity over fully converged kinetic models. In addition, the current PTM collection focuses on phosphorylation and methylation, and the sequence-to-dihedral prediction model was trained only on canonical peptides. Future versions will expand the resource to include additional PTMs, larger proteins, membrane proteins, ligand-bound systems, and other biologically relevant perturbations. Integration with enhanced sampling techniques and emerging AI-driven trajectory generation methods may further improve conformational coverage while maintaining physical consistency, extending pHaseMD4AI toward a more comprehensive phase-space resource for biomolecular simulation and machine learning.

## Conclusion

We present pHaseMD4AI, a molecular dynamics dataset that integrates globally equilibrated peptide simulations with protein-scale constant-pH MD simulations. The dataset combines canonical peptides, post-translationally modified peptides, peptide-CpHMD systems, and soluble proteins, providing synchronized coordinates, velocities, forces, protonation-state trajectories, and kinetic annotations where appropriate. By combining rigorous global-equilibrium characterization for peptide systems with broad pH-dependent sampling across folded proteins, pHaseMD4AI complements existing molecular dynamics resources and extends the range of physicochemical conditions represented in data-driven molecular modeling datasets. We anticipate that pHaseMD4AI will serve as a useful resource for developing and benchmarking molecular machine learning methods, including neural network potentials, generative models of biomolecular dynamics, and pH-aware modeling approaches, while also supporting broader studies of protein conformational dynamics, protonation-dependent behavior, and molecular simulation methodology.

## Availability and Community Contribution

The pHaseMD4AI dataset and associated processing scripts are available at https://isb.zju.edu.cn/md4ai/. To foster long-term community growth and data diversity, we have implemented a standardized upload interface. This allows other researchers to contribute additional peptide trajectories or pH-dependent protein data, which can be incorporated into the resource under branch-appropriate quality controls.

## Author contributions

Y.W. conceived and supervised the project; T.S. performed all simulations and analyzed all simulations together with J.H., Z.L., M.L., K.W., Y.Z. and Z.L. under the supervision of Y.W.; Y.G. built the web platform; Y.H. participated in discussions during the study and contributed to the development of the data; T.S. wrote the first draft of the manuscript with input from Y.W. All authors contributed to the writing of the manuscript.

## Supplementary materials

The Supporting Information is available free of charge at [Pending upon publication].

Kinetic consistency evaluation for HMR; algorithmic details of the sequence-based predictor; introduction to website design and functionality; comparison of Ramachandran landscapes across peptide systems; and data statistics for the protein-CpHMD branch.

## Data availability

The pHaseMD4AI post-processed data and associated post-processing scripts are available at https://isb.zju.edu.cn/md4ai/. The raw simulation data is available at https://ngdc.cncb.ac.cn/omix/release/OMIX018680.

## Acknowledgement

We thank Jiancheng Luo for assistance with CpHMD setup. We thank Hao Wu and Yan Wang for valuable discussions. Y.W. acknowledges the financial support of the Zhejiang Provincial National Science Foundation of China (No. LZ24C050003), the National Natural Science Foundation of China (No. 32371300), and the computational support of the Information Technology Center and State Key Lab of CAD&CG at Zhejiang University.

