## Supporting Information for "pHaseMD4AI: Phase-Space Dynamics Dataset with Chemical and pH Perturbations for Physically and Kinetically Consistent Biomolecular AI"

### 1 Ala2 kinetic consistency evaluation for HMR

#### 1.1 Basic experimental design

For two schemes of Ala2, we performed 5 $\mu$s simulations for each: (1) using the CHARMM36m force field with HMR, at a time step of 4 fs; and (2) using the CHARMM36m force field

[0]+ These authors contributed equally to this work.

without HMR, at a time step of 2 fs. We considered the distances between all backbone heavy atoms as features. To ensure consistency of comparison, we performed dimensionality reduction using TICA and subsequent k-means clustering. We then constructed Markov state models (MSMs) and the Robust Perron-cluster cluster analysis (PCCA+) for each scheme.

##### 1.2 Detailed Parameters

We obtained a total of 55 distance pairs between backbone heavy atoms. We set the feature extraction frequency to 10 ps/frame, the TICA lag time to 10 ps, k-means to cluster the initial trajectories into 100 classes on the first 10 dimensions of TIC, the MSM lag time to 50 ps, and the macrostate classes to 4. We calculated the probability of the macrostate and the mean first passage time (MFPT) between macrostates.

##### 1.3 Results

The results showed that the metastable state distributions and interstate transition rates were almost identical for schemes with and without HMR enabled (Fig. S). This validation confirms that using HMR with a 4 fs time step does not alter the fundamental dynamics or free energy landscape of the peptide system.

#### 2 Prediction of Dihedral Angle Distributions from Sequence

##### 2.1 Algorithm design and target representation

The goal of this application was to predict the equilibrium dihedral distributions of non-terminal peptide residues directly from sequence. We constructed it from the standard-peptide core branch, using the 20 tripeptides and 400 tetrapeptides in the globally equilibrated standard-peptide subset, for a total of 420 systems. Ala capping residues were treated as fixed context, and only the non-terminal residues were used as prediction targets.

For every target residue, the backbone Ramachandran distribution was represented by a periodic $72 \times 72$ histogram over $(\phi, \psi)$, and each available side-chain torsion $\chi_1$–$\chi_4$ was represented by a periodic 72-bin one-dimensional histogram. This makes the model output explicit at the level of individual dihedral degrees of freedom, including residue-wise backbone and side-chain torsions. All angles were wrapped to $[-\pi, \pi)$ before histogramming, and each histogram was normalized to a probability distribution. Residues without a given $\chi$ torsion were masked rather than assigned artificial zero-valued targets. Free-energy landscapes were not predicted directly; when needed for visualization, they were derived post hoc from the probability distributions according to

$$F = -k_B T \ln p + C, \tag{1}$$

where $C$ shifts the minimum free energy to zero.

#### 2.2 Train/validation/test split

For the cross-length generalization setting described in the main text, all 20 tripeptides were used for model development and all 400 tetrapeptides were reserved as an external test set.

#### 2.3 Stage-1 landscape VAE

The first stage compresses residue-resolved dihedral distributions into a compact latent space. The model is a slot-aware landscape variational autoencoder[1] with separate encoders for backbone and side-chain distributions. Ramachandran maps were encoded with circular 2D convolutional blocks, and $\chi$ histograms were encoded with circular 1D convolutional blocks so that the periodic boundary at $\pm\pi$ was handled explicitly. The encoded residue-wise features were then combined into two latent levels: a residue-specific slot latent of dimension 8 and a peptide-level global latent of dimension 16. The hidden dimension used in the shared multilayer perceptrons was 256, and the model allowed up to two variable positions per peptide.

The stage-1 model was trained with Adam using learning rate $10^{-3}$, batch size 32, and 200 epochs. The loss was the sum of Ramachandran cross-entropy, $\chi$ cross-entropy, and Kullback–Leibler penalties on the global and slot latent variables:

$$\mathcal{L}_{\text{VAE}} = \mathcal{L}_{\phi\psi}^{\text{CE}} + \mathcal{L}_{\chi}^{\text{CE}} + \beta_g \mathcal{L}_{\text{KL}}^{\text{global}} + \beta_s \mathcal{L}_{\text{KL}}^{\text{slot}}, \tag{2}$$

with $\beta_g = \beta_s = 10^{-3}$.

#### 2.4 Stage-2 sequence-to-latent model

The second stage predicts the VAE latent variables directly from sequence. We used a frozen ESM2_t6_8M_UR50D checkpoint,[2] corresponding to the 6-layer, ~8M-parameter member of the ESM2 family, to provide two types of sequence representation: (i) a full-sequence embedding obtained by mean-pooling residue token representations over the sequence and used by the peptide-level branch and (ii) residue-token embeddings extracted at the variable positions and used by the slot-level branch. These learned sequence features were combined with explicit residue descriptors.

For each target residue, the slot branch received the concatenation of the residue-level ESM embedding, a 20-dimensional amino-acid one-hot vector, an 11-dimensional physicochemical feature vector, an 8-dimensional slot-position embedding, and a binary slot-validity mask. The 11 explicit physicochemical features were hydrophobicity on the Kyte–Doolittle scale, number of side-chain heavy atoms, number of side-chain $\chi$ torsions, formal charge sign $(-1/0/+1)$, aromaticity, branching, polarity, sulfur content, hydrogen-bonding capacity, glycine identity, and proline identity. For the peptide-level branch, the global explicit feature vector had dimension 22 and consisted of the mean of the slot property vectors concatenated with their absolute difference. Slot and global branches were each modeled by two-layer multilayer perceptrons with hidden dimension 256. The stage-2 mapper was trained for 100 epochs with Adam, learning rate $10^{-3}$, and batch size 16.

The stage-2 loss combined latent matching and distribution reconstruction:

$$\mathcal{L}_{\text{mapper}} = \mathcal{L}_{\text{latent}} + \mathcal{L}_{\text{recon}}, \tag{3}$$

where $\mathcal{L}_{\text{latent}}$ is the mean squared error between predicted and ground-truth global/slot latent means, and $\mathcal{L}_{\text{recon}}$ is the same distribution cross-entropy used in stage 1 after passing the predicted latent variables through the frozen VAE decoder.

#### 2.5 Evaluation metrics and results

Performance was measured on the held-out tetrapeptide test set. Besides the training losses, we used Jensen–Shannon similarity[3] to quantify the agreement between predicted and ground-truth probability distributions. For two normalized distributions $p$ and $q$, this similarity is defined as

$$\text{JS similarity}(p, q) = 1 - \frac{\text{JS}(p, q)}{\ln 2}, \tag{4}$$

so that a value of 1 indicates perfect agreement. Ramachandran and side-chain distributions were summarized separately, averaging only over valid target residues and valid $\chi$ torsions. On the 400-system tetrapeptide test set, this corresponds to 800 residue-wise Ramachandran maps and 1560 valid $\chi$ curves.

The stage-1 landscape VAE reconstructed the held-out tetrapeptide distributions with good fidelity under this cross-length setting. On the test set, the mean JS similarity was 0.9478 for Ramachandran distributions and 0.9609 for side-chain $\chi$ distributions.

The stage-2 sequence-to-latent model achieved similar but slightly lower reconstruction quality on the same tetrapeptide test set. The mean JS similarity was 0.9481 for Ramachandran distributions and 0.9618 for side-chain $\chi$ distributions, with a latent-space mean squared error of 0.0364. These values indicate that the frozen-decoder route preserves most of the landscape representation learned in stage 1 even when the sequence-conditioned model is evaluated under a tripeptide-to-tetrapeptide generalization setting.

### 3 Overall design logic of the website

pHaseMD4AI is a web portal for browsing, viewing, and downloading molecular dynamics datasets. The website follows the usual order in which users approach the data: a general overview first, then system-level search and filtering, and finally entry-specific visualization and analysis. A user may start from the homepage, move to the Search/Browse page to identify a peptide or protein of interest, check the Residues page when sequence annotations or modification labels need clarification, and then open the corresponding Workbench page. The Workbench page brings the structure viewer, trajectory-related plots, analysis panels, and download links together in one place. Separate pages are also provided for dataset statistics, workflow documentation, data upload, and archive access.

The current release includes tripeptides, tetrapeptides, proteins, post-translationally modified variants, and constant-pH molecular dynamics cases. The main website components described in the appendix are therefore: (1) homepage/platform overview, (2) Search/Browse page, (3) Residues page, (4) system-specific Workbench page, (5) Statistics page, (6) Documentation page, (7) Upload page, and (8) Archive page. Within the Workbench page, the main elements are the system summary and method switcher, trajectory viewer, potential-energy plot, free-energy surface and coarse-grained state panel, convergence diagnostics, method comparison, and download section.

### 4 Module-by-module description

#### Module 1. Homepage

The homepage gives a general overview of pHasepHaseMD4AI. It introduces the website as a phase-space molecular dynamics dataset covering systems with chemical modification and pH-dependent perturbation conditions. A central search bar supports queries by sequence or protein identifier. It links directly to the corresponding analysis page. The homepage also displays summary numbers for tripeptides, tetrapeptides, proteins, and total entries. The navigation bar leads to the main sections of the website, including Search/Browse, Residues, Statistics, Documentation, Upload, and Archive.

#### Module 2. Search/Browse page

The Search/Browse page is used to locate entries in the dataset. Searches can be performed by sequence, PDB identifier, modification label, or protonation-state annotation. Filters are available for system class, post-translational modification category, and CpHMD annotation. Each table entry lists the system sequence or identifier, sequence length, modification status, pH-related condition, available simulation methods, and a structure preview. The View link opens the corresponding Workbench page.

#### Module 3. Residues page

The Residues page lists the residue types used in the dataset. It includes standard amino acids together with modified and protonation-dependent residue variants. Searches can be made by residue name, one-letter code, PTM label, phosphorylation label, or CpHMD-related notation. Each residue entry includes a schematic structure and a short annotation. The residue cards also link back to the Search/Browse page with the relevant query applied. This makes it easier to find systems containing that residue or modification.

#### Module 4. Workbench page

After selecting a system, the user enters the Workbench page. This is the main entry-specific page of the website. It combines metadata, structure visualization, trajectory display, analysis results, and downloadable files in a single layout. For each system, the page is divided into several panels corresponding to different parts of the simulation output.

##### A. System summary and method switching

At the top of the Workbench page, basic information about the selected system is shown, including sequence length, available simulation methods, equilibrium status, and update time. This section also includes the method switcher. It toggles between Unbiased MD and Adaptive sampling views for the same entry. For constant-pH cases, protonation-state conditions may also appear at the entry level.

##### B. Trajectory viewer and metadata panel

The main visualization area contains an interactive trajectory viewer based on MDsrv/NGL for streaming PDB and XTC files. Users can play the trajectory, step through frames, and reset the molecular view. Vector overlays such as force and velocity are also available, together with controls for vector length, cutoff settings, and atom selection.

A metadata panel beside the viewer summarizes the main properties of the current system, including sequence length, available methods, force field, temperature, and equilibrium status.

##### C. Potential-energy inspection

A potential-energy panel is shown for the currently selected simulation method. The energy plot is linked to the trajectory viewer. The corresponding structural frame can therefore be inspected together with the energy value. This makes it possible to compare conformational changes with the energy profile along the trajectory.

##### D. Free-energy surface and coarse-grained states

Another panel shows the free-energy surface together with the associated coarse-grained states. This section presents the sampled conformational landscape in a reduced coordinate space and displays the state-based organization of the simulation results. Users can inspect the main regions of the landscape and their associated conformational states.

##### E. Convergence diagnostics and method comparison

The Workbench page also includes a convergence-diagnostics panel and a method-comparison panel. The convergence panel shows diagnostic figures used to assess sampling behavior for the current system. The comparison panel places results from different simulation strategies, especially Unbiased MD and Adaptive sampling, side by side for the same entry.

##### F. Downloads and data access

The final section of the Workbench page lists the downloadable files associated with the selected system and simulation method. These files are grouped into categories such as trajectory-related files, feature-related files, MSM-related files, figures, and bundled archives containing the full set of outputs.

#### Module 5. Statistics page

The Statistics page summarizes the dataset at the global level. It reports the total number of entries, the total simulation time represented in the collection, and the approximate size of the uncompressed data. It also includes plots showing the distribution of system classes, the distribution of trajectory lengths, and protein-related statistics such as residue counts and titratable-site counts.

##### Module 6. Documentation page

The Documentation page outlines the simulation and analysis workflow used for the dataset. It describes the workflow in two parts: sampling and post-processing. The sampling section covers conventional unbiased molecular dynamics and adaptive sampling. The post-processing section describes feature extraction, dimensionality reduction, clustering, Markov-state-model construction, coarse graining, and the calculation of free-energy and kinetic quantities. The page also lists the main workflow scripts used to reproduce or extend the pipeline.

##### Module 7. Upload page

The Upload page is used for dataset submission. It accepts peptide or protein molecular dynamics datasets as compressed archives together with basic metadata, including protein name, file description, and contact email.

##### Module 8. Archive page

The Archive page gives access to the large-scale raw dataset associated with pHasepHaseMD4AI. It points to the external bulk archive hosted through the National Center for Bioinformatics, China.

#### 5. Supplementary Figures

(2) Lin, Z.; Akin, H.; Rao, R.; Hie, B.; Zhu, Z.; Lu, W.; Dos Santos Costa, A.; Fazel-

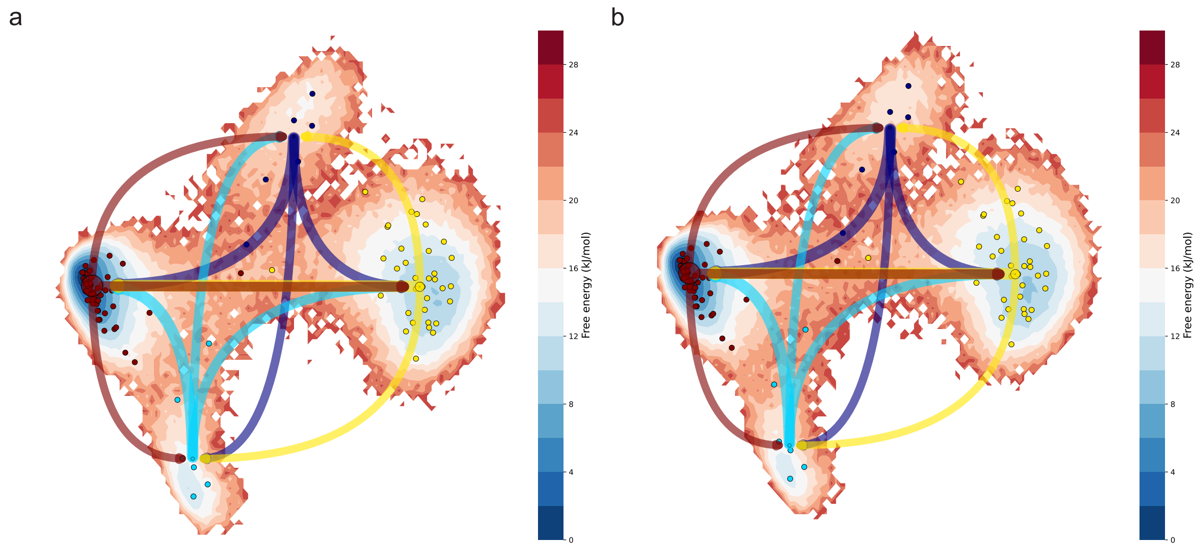


**Figure S1:** HMR Kinetic Validation. Markov state models of the alanine dipeptide show identical metastable states and transition kinetics with HMR at a 4 fs timestep and without HMR at a 2 fs timestep. Projections of HMR (left) and no HMR (right) onto the first two dimensions of tICs: free energy topography, macrostate population, and transition rate. Each small circle represents a microstate. Each large circle represents a macrostate, including the microstates of the same color, and its size is proportional to the population of that state. The lines represent the transition rate between every two macrostates, and the line thickness is proportional to $\log_2(1/\mathrm{MFPT}_{ij})$.

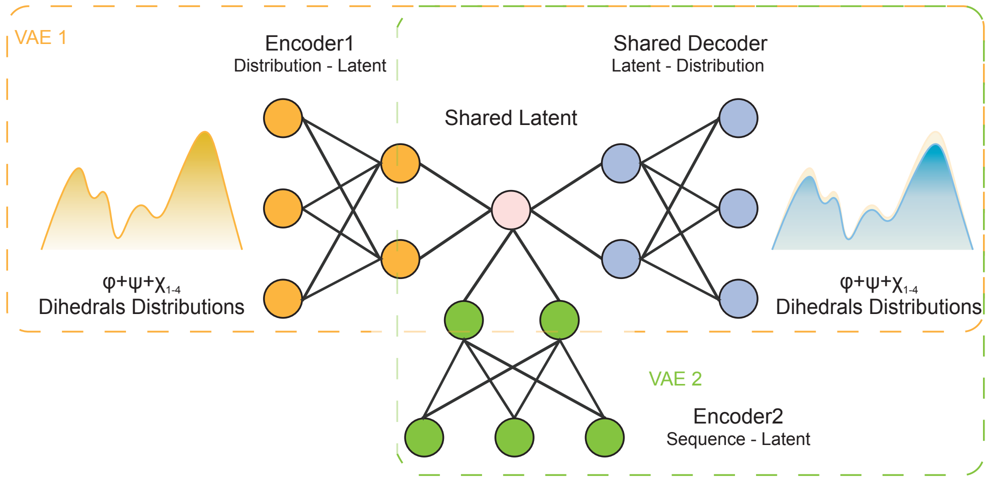


**Figure S2:** Schematic overview of the two-stage prediction framework, in which a distribution variational autoencoder learns latent representations of residue-resolved dihedral distributions and a sequence-conditioned predictor maps sequence features to the latent space for distribution reconstruction.

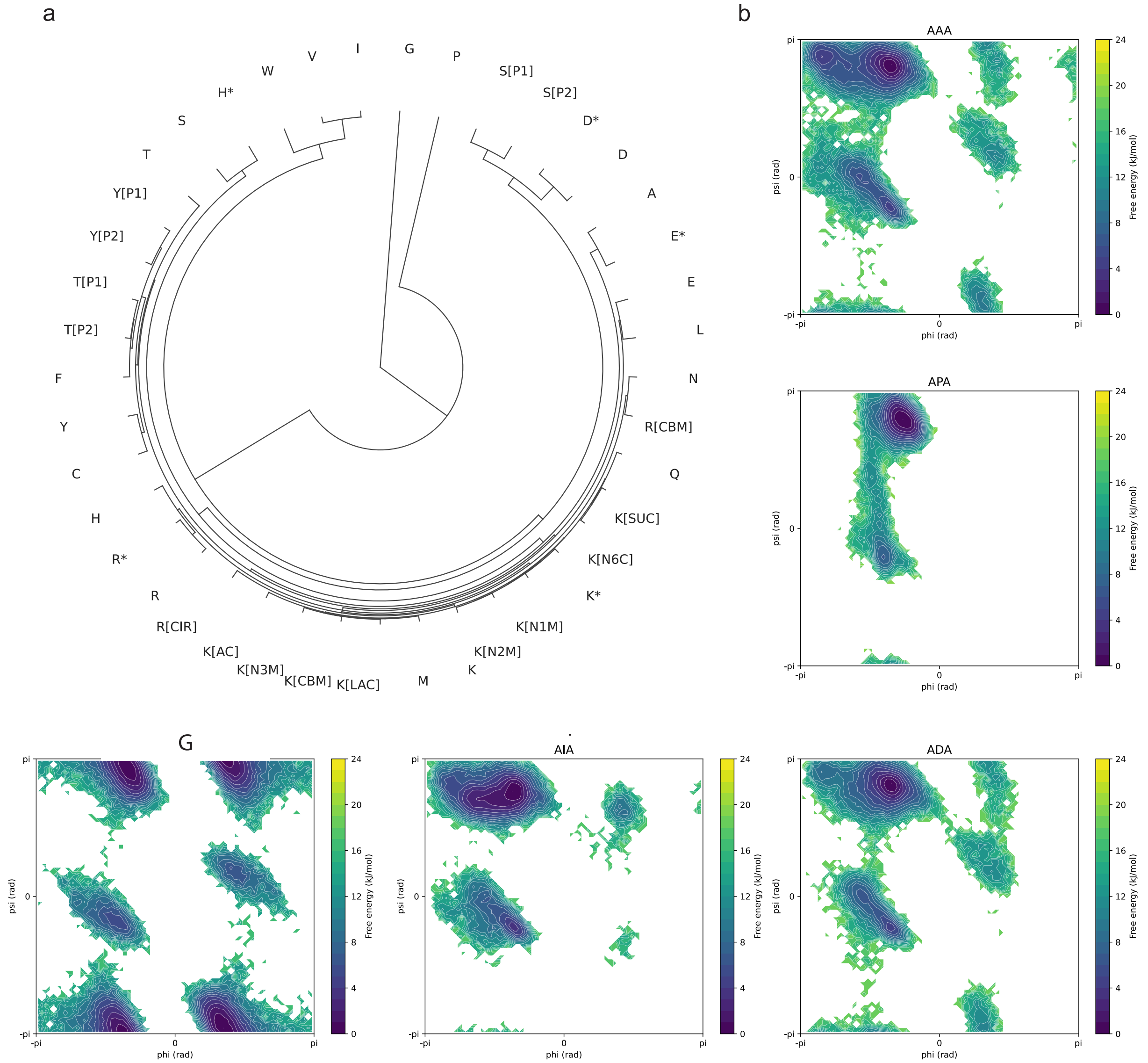


**Figure S3:** Hierarchical clustering of tripeptides (a) and $\phi - \psi$ 2-dimensional FES of representative residues (b).

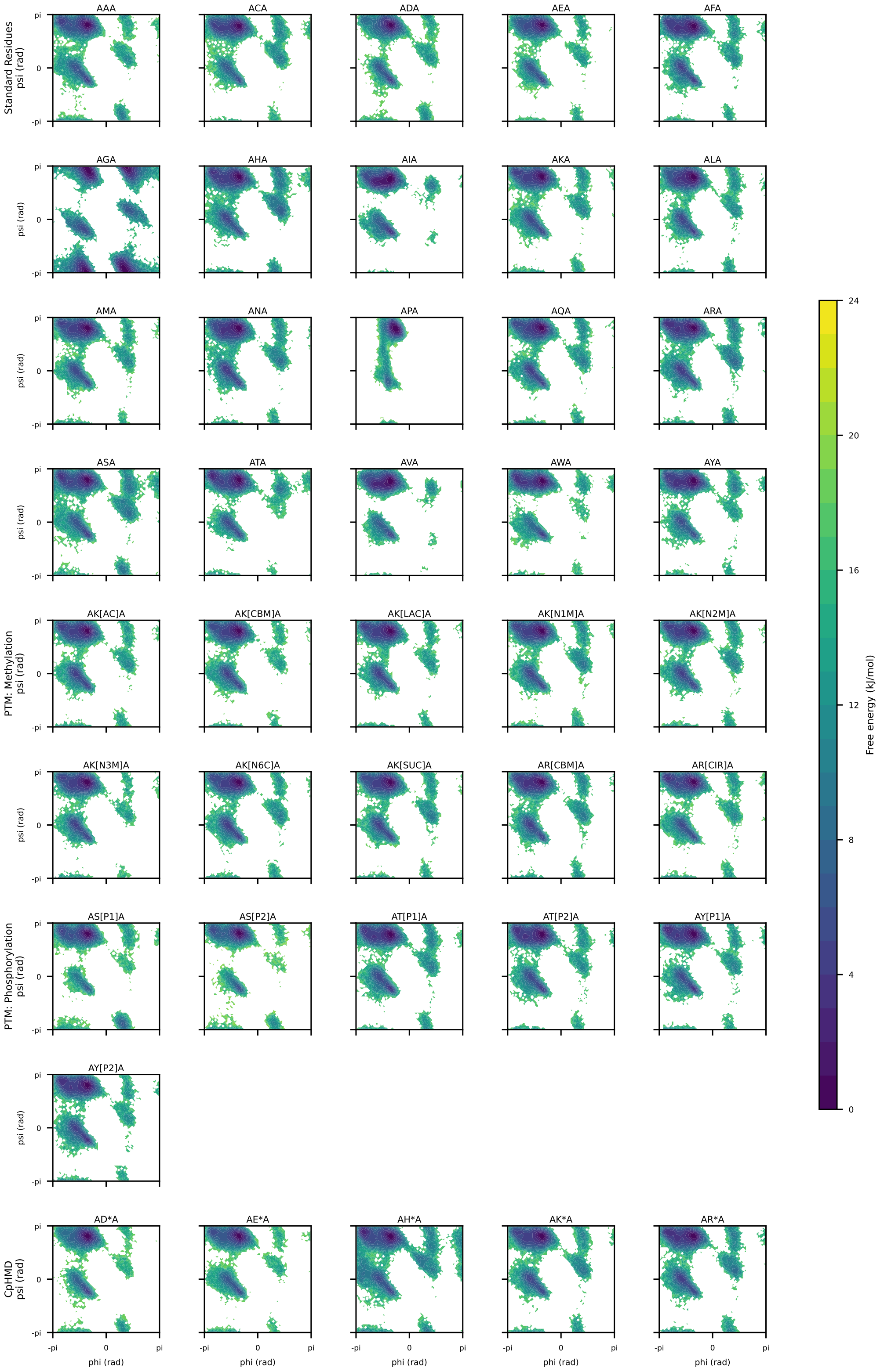


**Figure S4:** Tripeptide Ramachandran Free-Energy Landscapes across Standard, PTM, and CpHMD Systems.

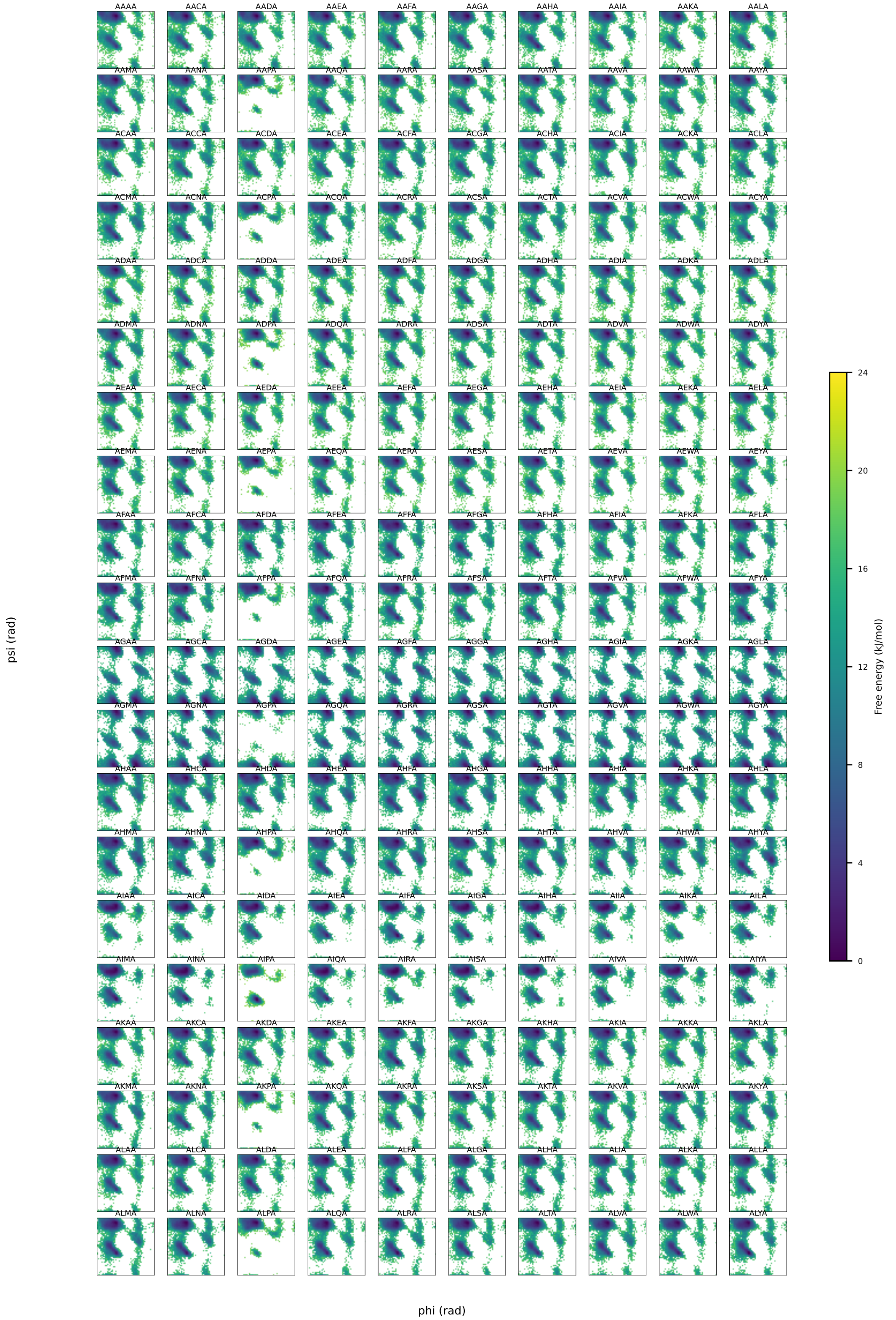

**Figure S5:** Standard Tetrapeptide $\phi_1 - \psi_1$ Free-Energy Landscapes (1-200/400).

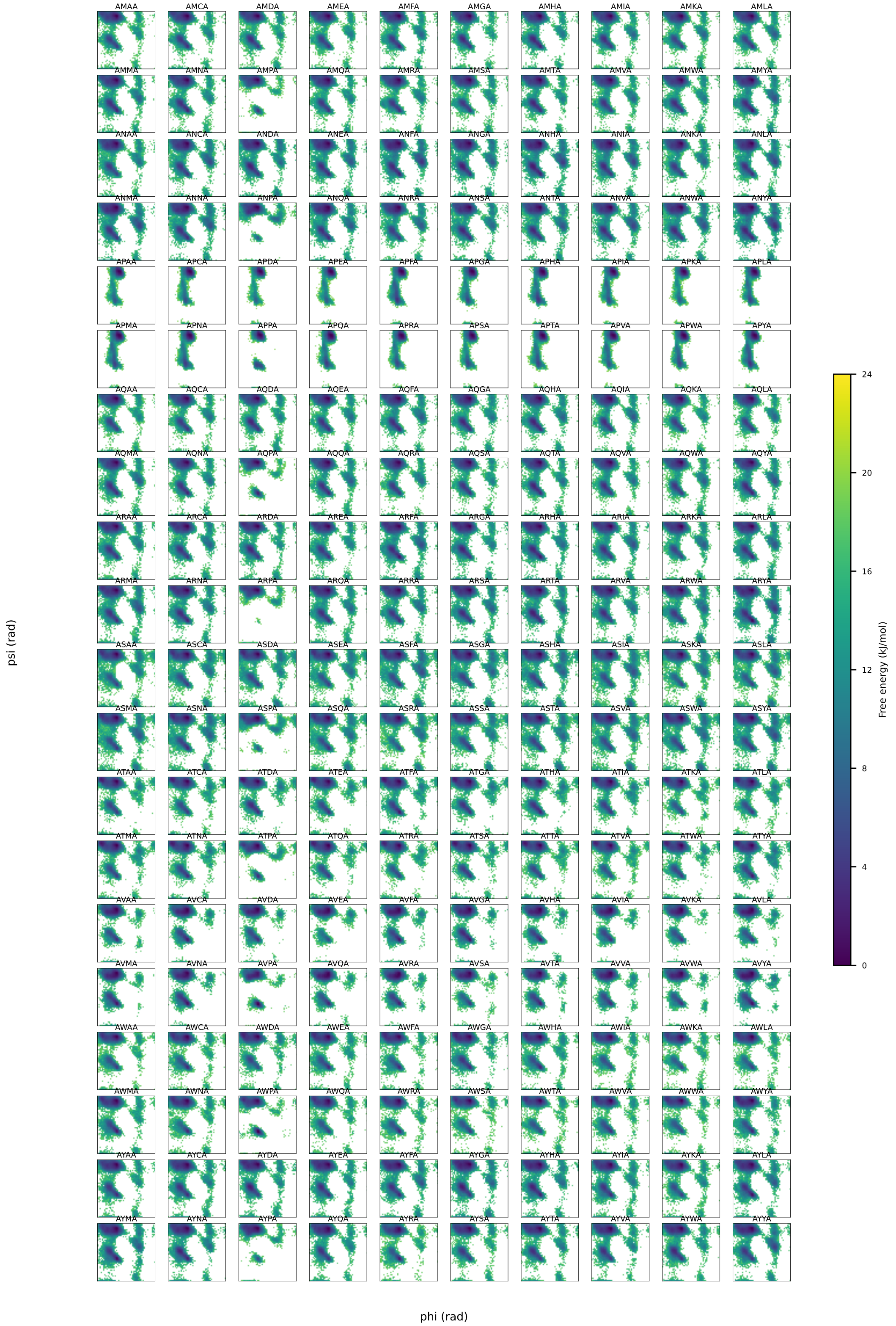


**Figure S6:** Standard Tetrapeptide $\phi_1 - \psi_1$ Free-Energy Landscapes (201-400/400).

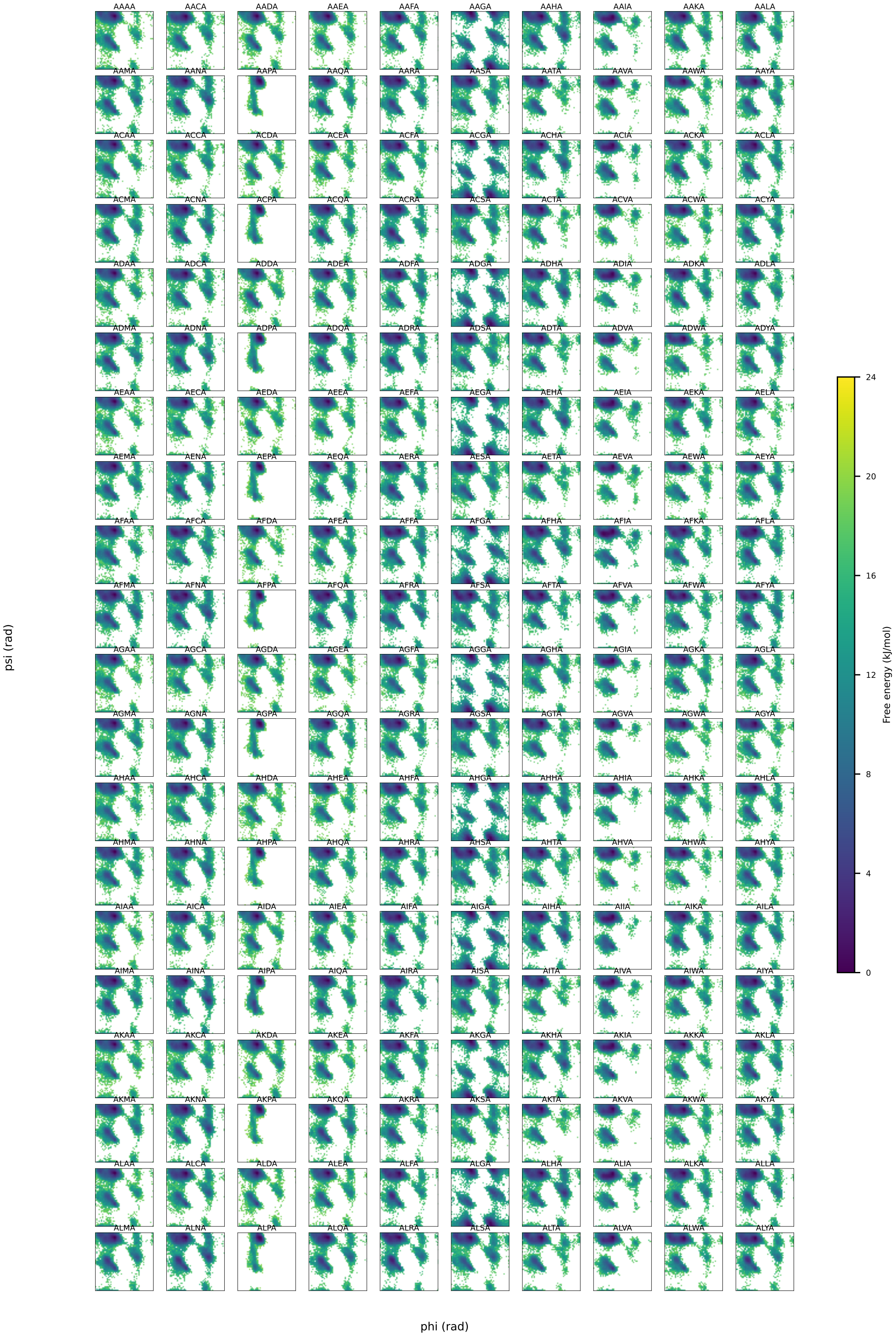


**Figure S7:** Standard Tetrapeptide $\phi_2 - \psi_2$ Free-Energy Landscapes (1/200).

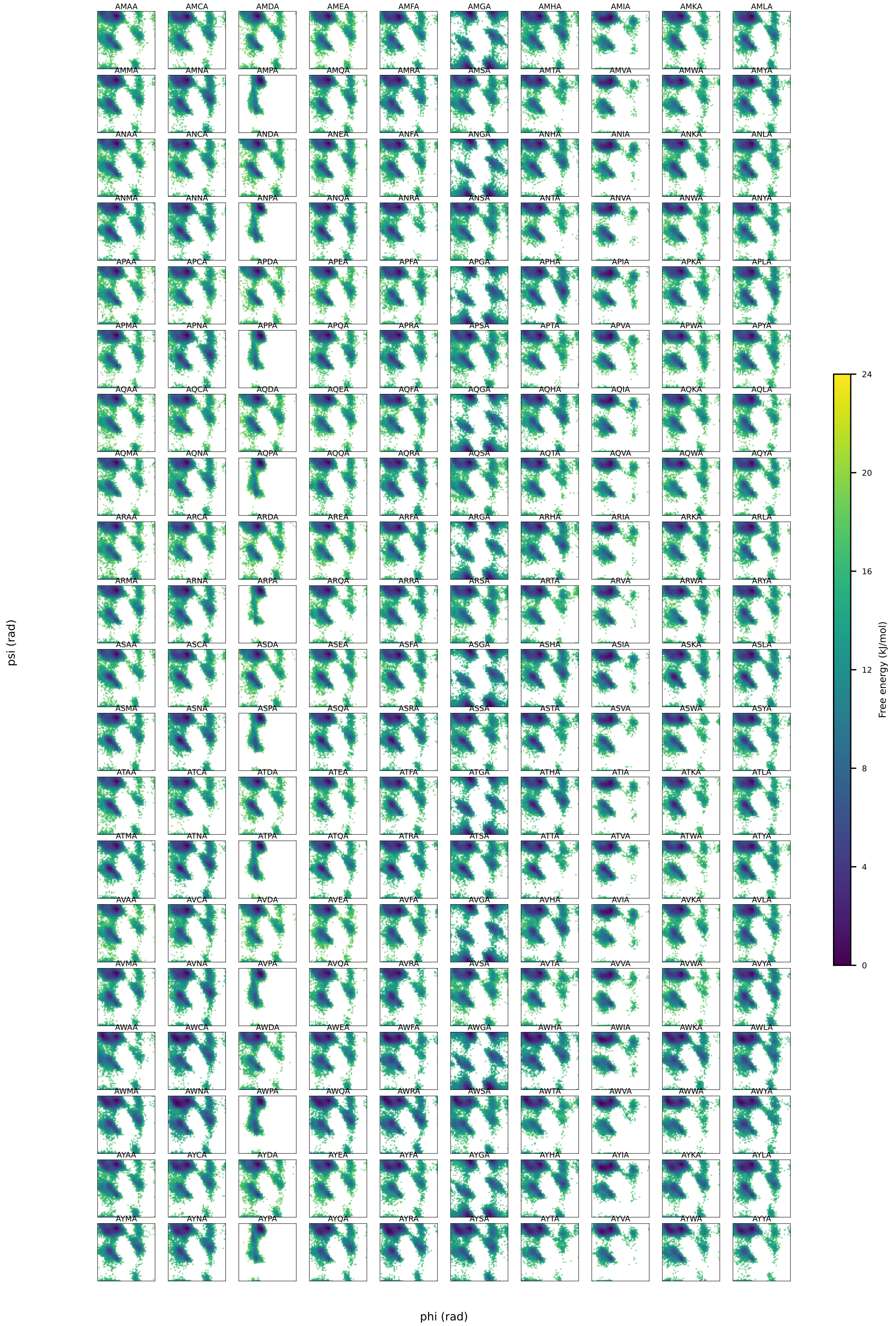


**Figure S8:** Standard Tetrapeptide $\phi_2 - \psi_2$ Free-Energy Landscapes (201-400/400).

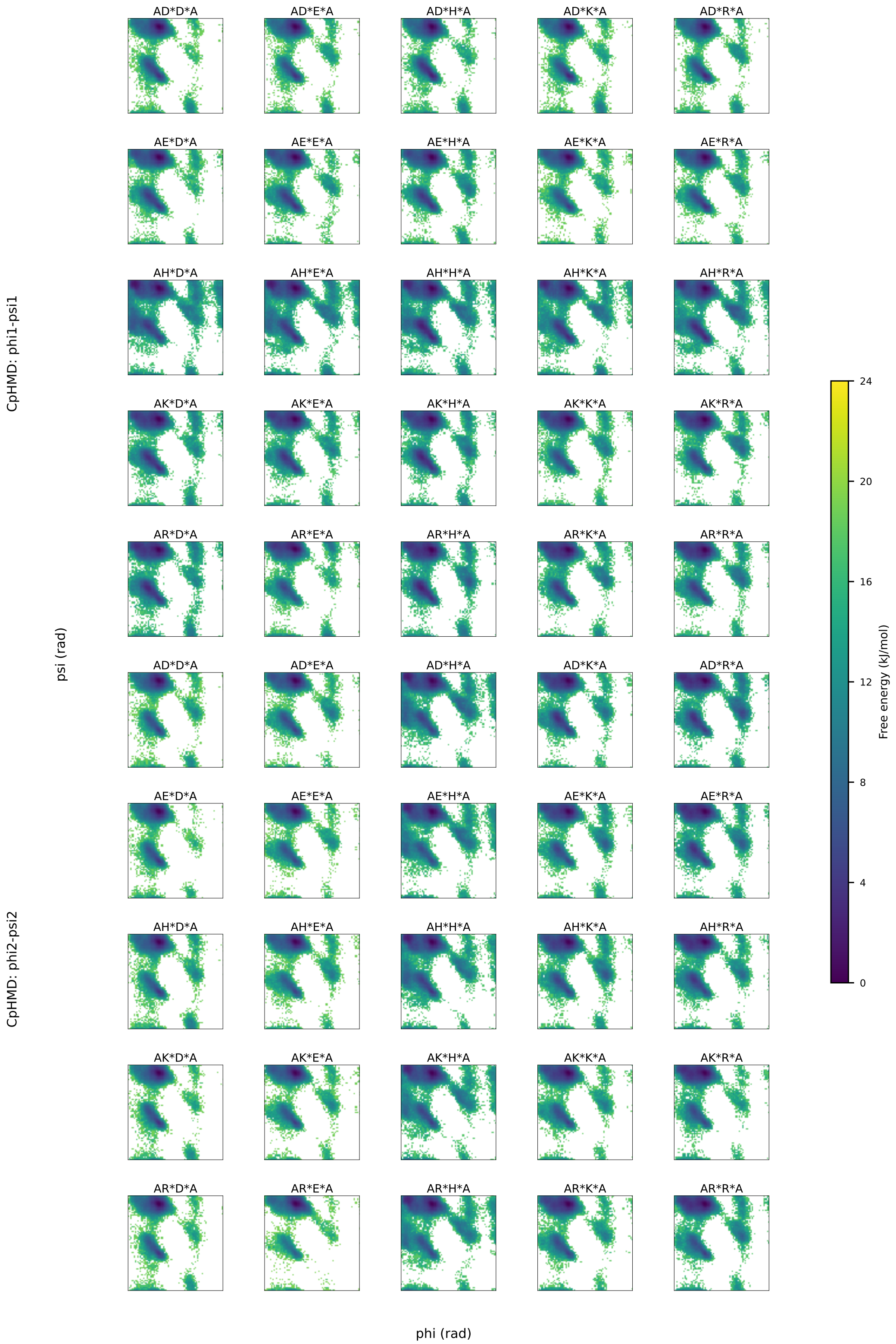


**Figure S9:** CpHMD Tetrapeptide $\phi_1 - \psi_1$ and $\phi_2 - \psi_2$ Free-Energy Landscapes.

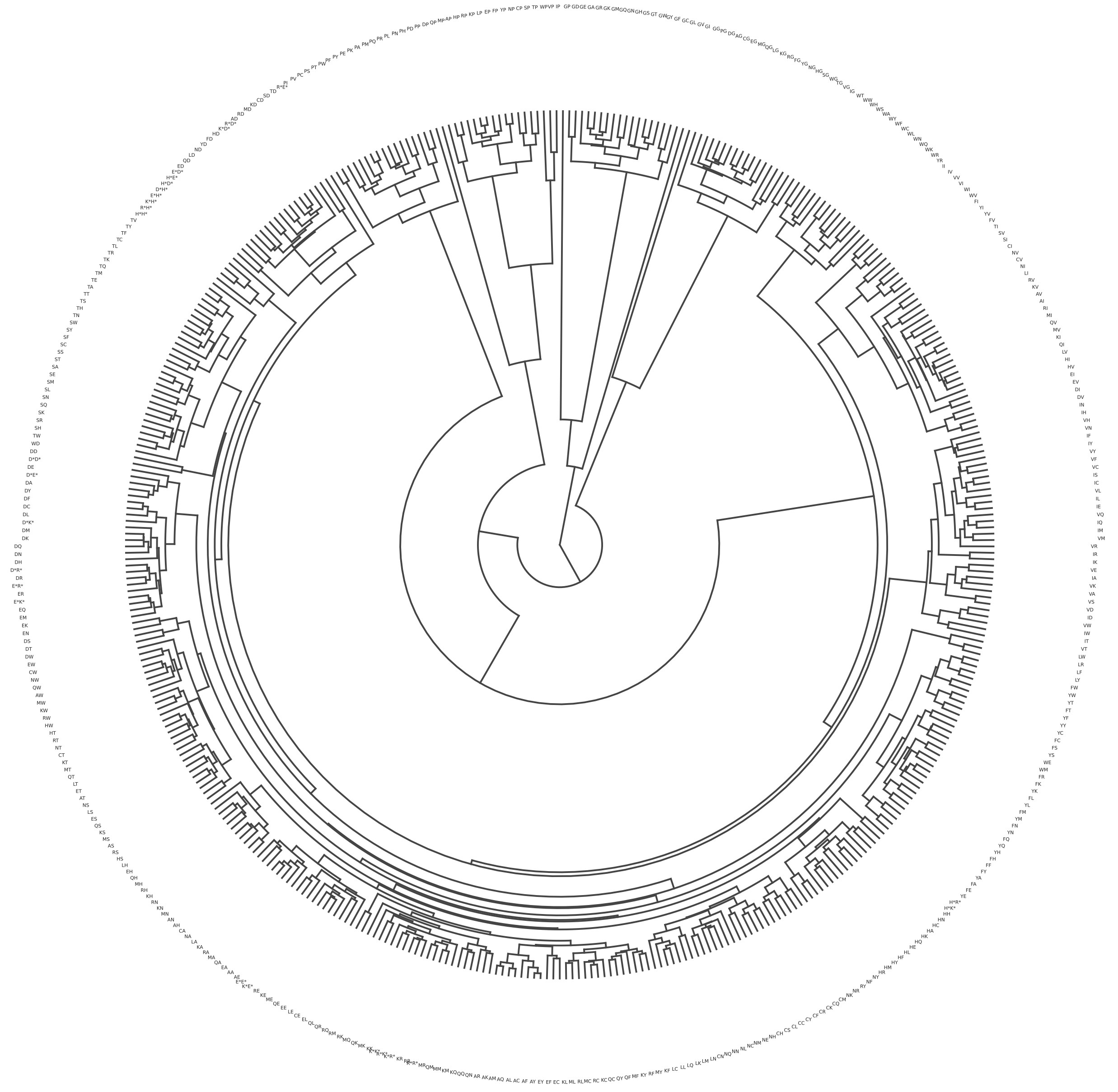

**Figure S10:** Hierarchical clustering of tetrapeptides based on cross-system Ramachandran similarity analysis.

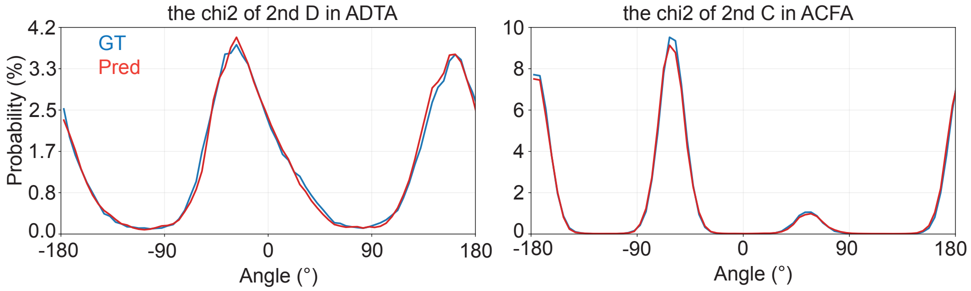


**Figure S11:** Representative comparisons between predicted and ground-truth 1D side-chain dihedral probability distributions for test-set examples, illustrating that the model captures the distinct torsional population patterns.

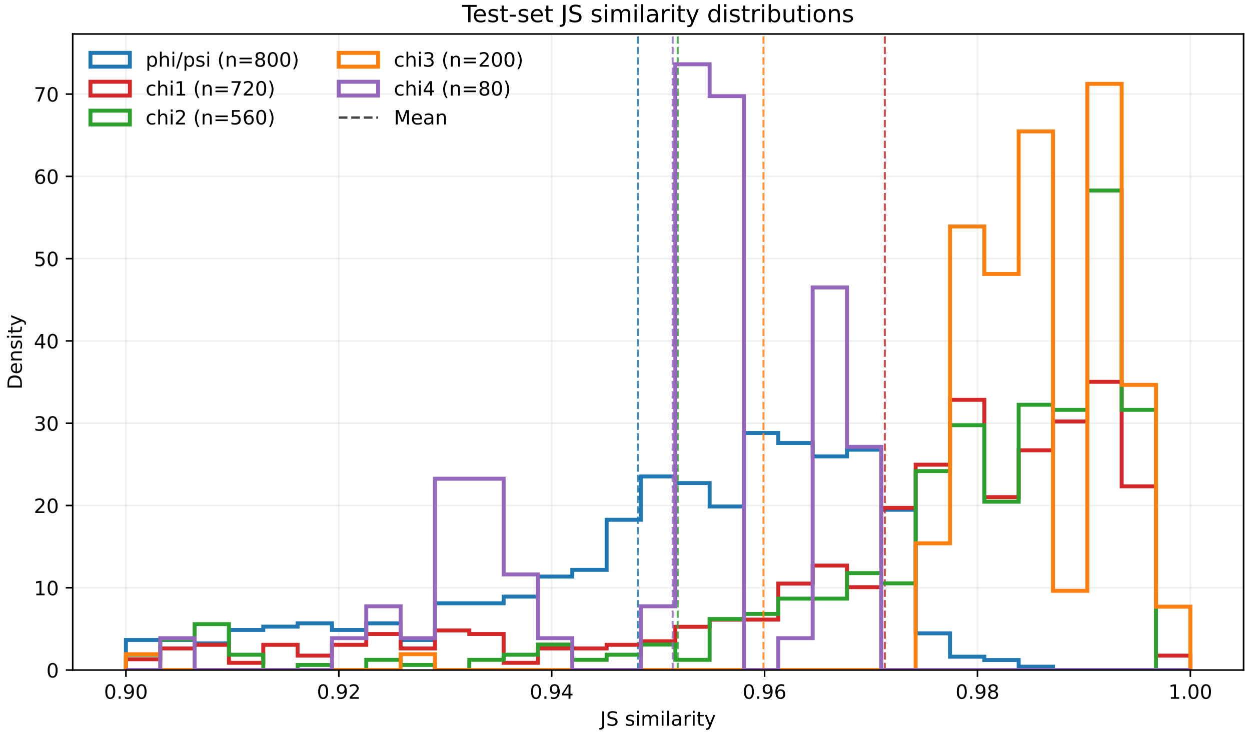


**Figure S12:** Jensen-Shannon similarity distributions on the tetrapeptide test set.

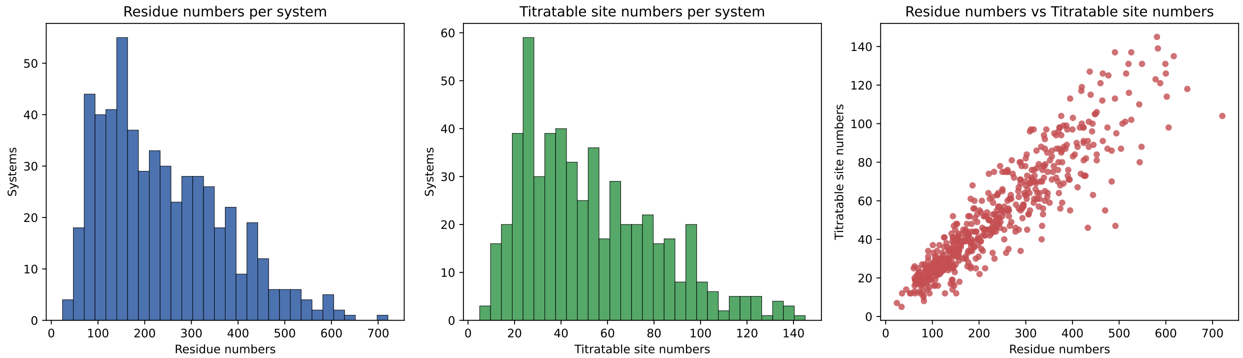


**Figure S13:** Histograms of residue numbers, titratable site numbers, and their relationship across Protein-CpHMD branch.

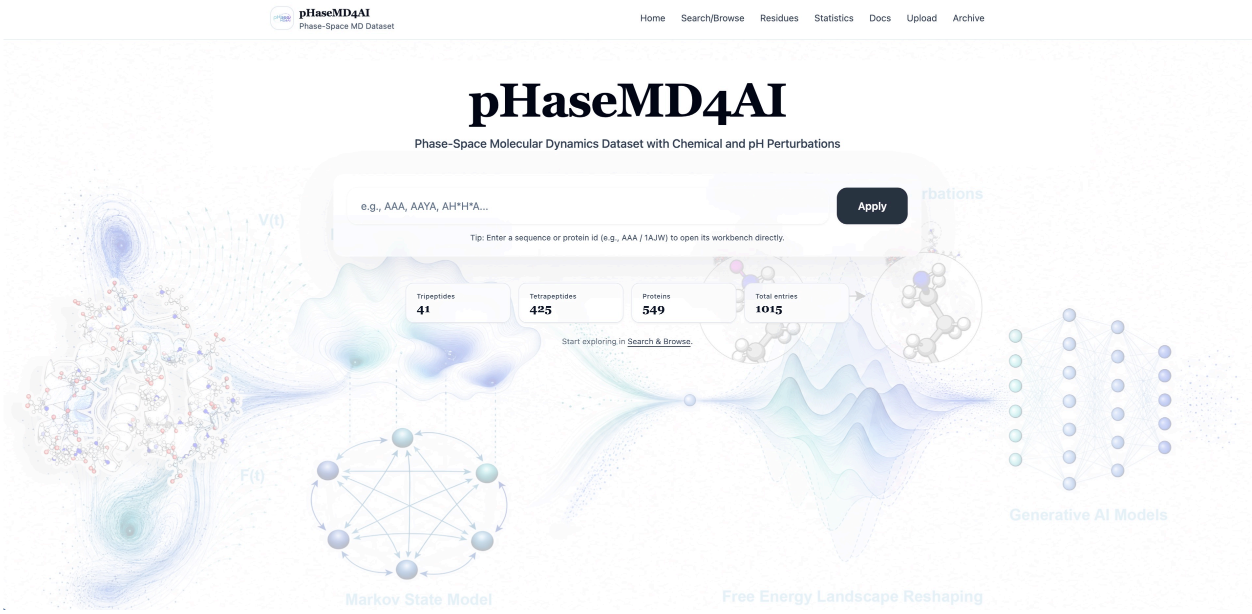


**Figure S14:** Homepage of the pHaseMD4AI web platform.

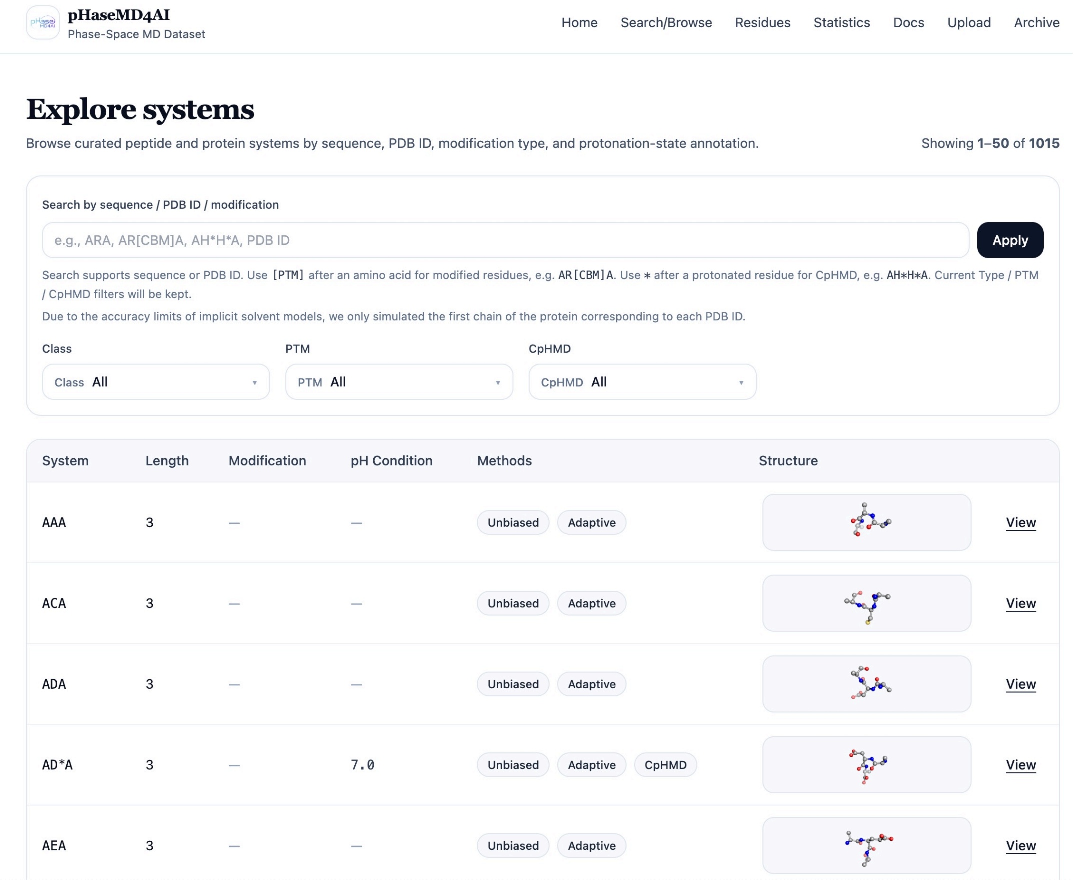


**Figure S15:** Search/Browse page of the pHaseMD4AI web platform.

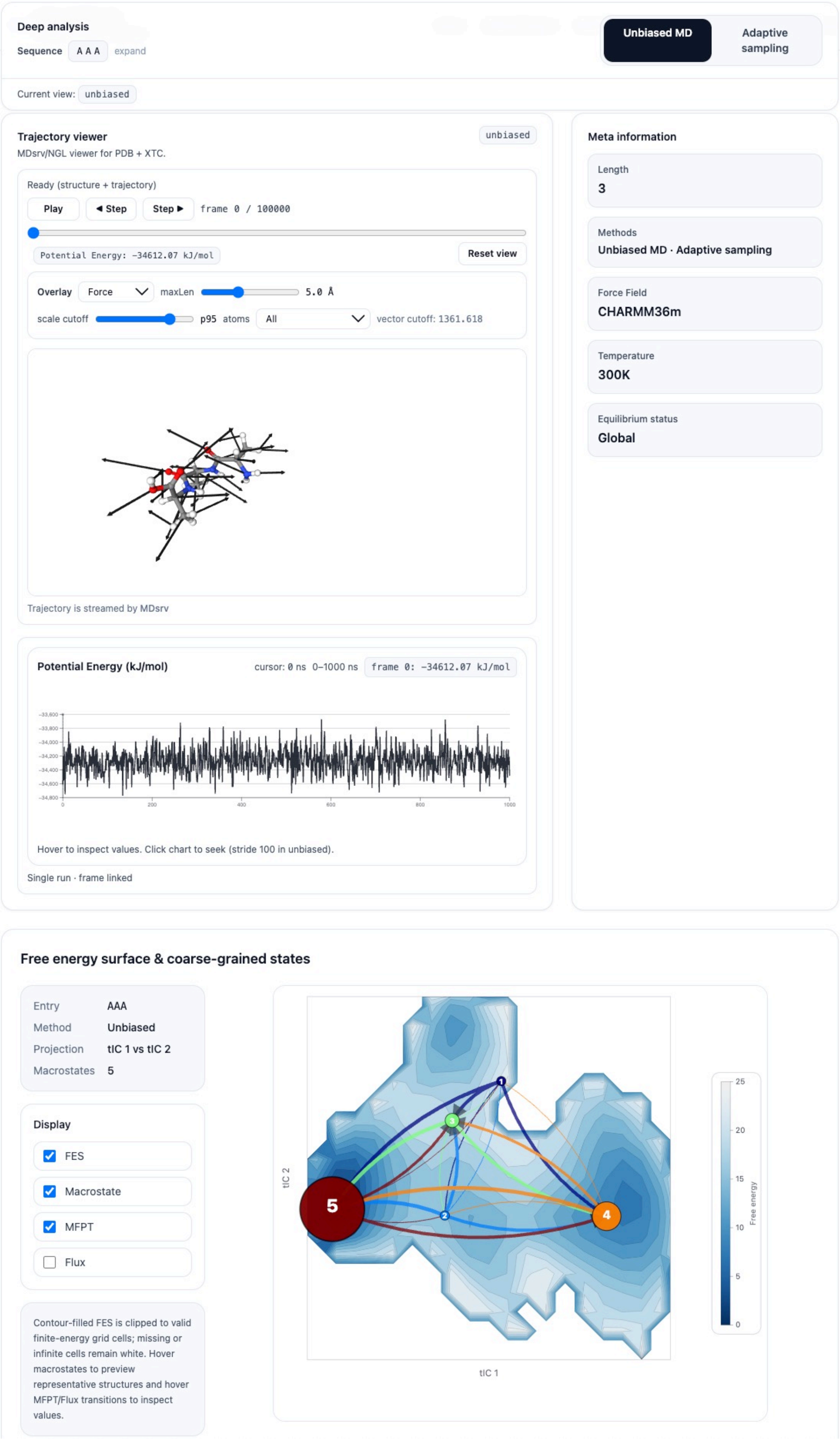


**Figure S16:** Workbench page of the pHaseMD4AI web platform.

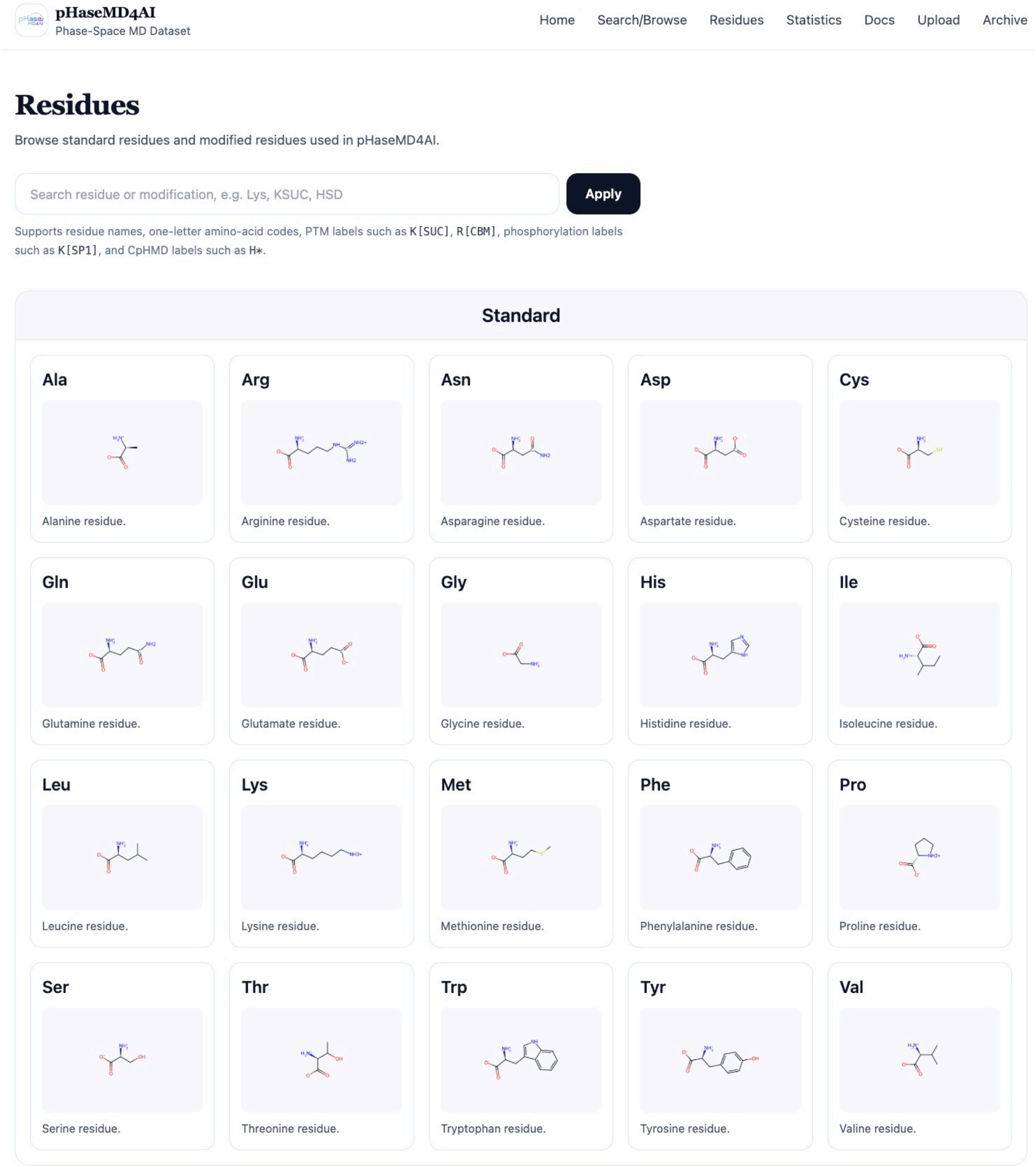


**Figure S17:** Residues page of the pHaseMD4AI web platform.

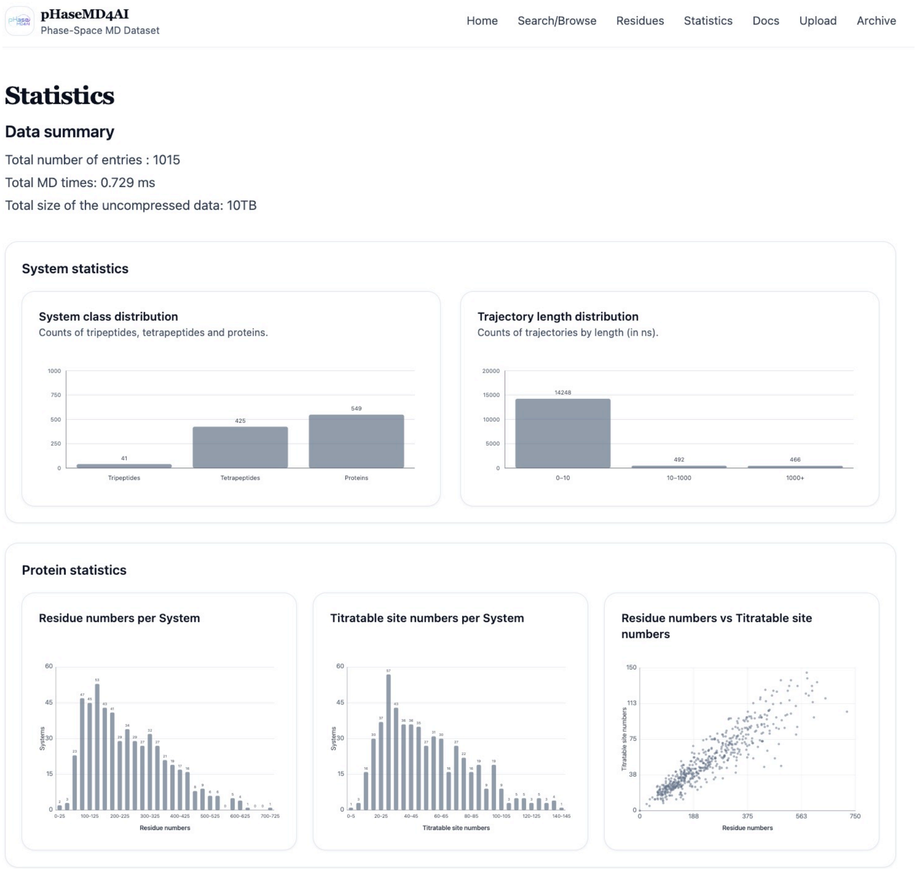


**Figure S18:** Statistics page of the pHaseMD4AI web platform.

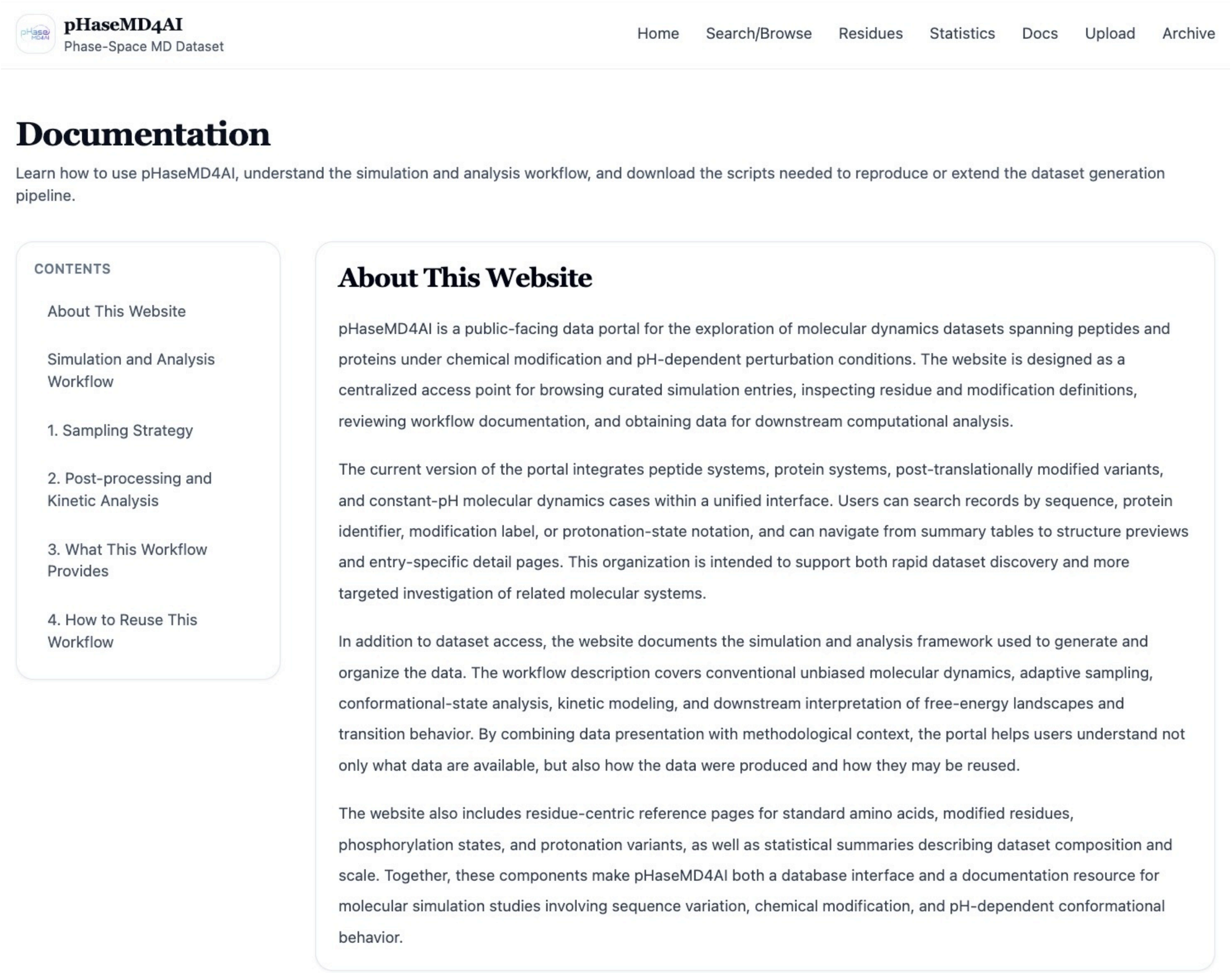

pHaseMD4AI
Phase-Space MD Dataset

Home Search/Browse Residues Statistics Docs Upload Archive

### Documentation

Learn how to use pHaseMD4AI, understand the simulation and analysis workflow, and download the scripts needed to reproduce or extend the dataset generation pipeline.

CONTENTS


About This Website
Simulation and Analysis Workflow
1. Sampling Strategy
2. Post-processing and Kinetic Analysis
3. What This Workflow Provides
4. How to Reuse This Workflow


#### About This Website

pHaseMD4AI is a public-facing data portal for the exploration of molecular dynamics datasets spanning peptides and proteins under chemical modification and pH-dependent perturbation conditions. The website is designed as a centralized access point for browsing curated simulation entries, inspecting residue and modification definitions, reviewing workflow documentation, and obtaining data for downstream computational analysis.

The current version of the portal integrates peptide systems, protein systems, post-translationally modified variants, and constant-pH molecular dynamics cases within a unified interface. Users can search records by sequence, protein identifier, modification label, or protonation-state notation, and can navigate from summary tables to structure previews and entry-specific detail pages. This organization is intended to support both rapid dataset discovery and more targeted investigation of related molecular systems.

In addition to dataset access, the website documents the simulation and analysis framework used to generate and organize the data. The workflow description covers conventional unbiased molecular dynamics, adaptive sampling, conformational-state analysis, kinetic modeling, and downstream interpretation of free-energy landscapes and transition behavior. By combining data presentation with methodological context, the portal helps users understand not only what data are available, but also how the data were produced and how they may be reused.

The website also includes residue-centric reference pages for standard amino acids, modified residues, phosphorylation states, and protonation variants, as well as statistical summaries describing dataset composition and scale. Together, these components make pHaseMD4AI both a database interface and a documentation resource for molecular simulation studies involving sequence variation, chemical modification, and pH-dependent conformational behavior.

**Figure S19:** Docs page of the pHaseMD4AI web platform.

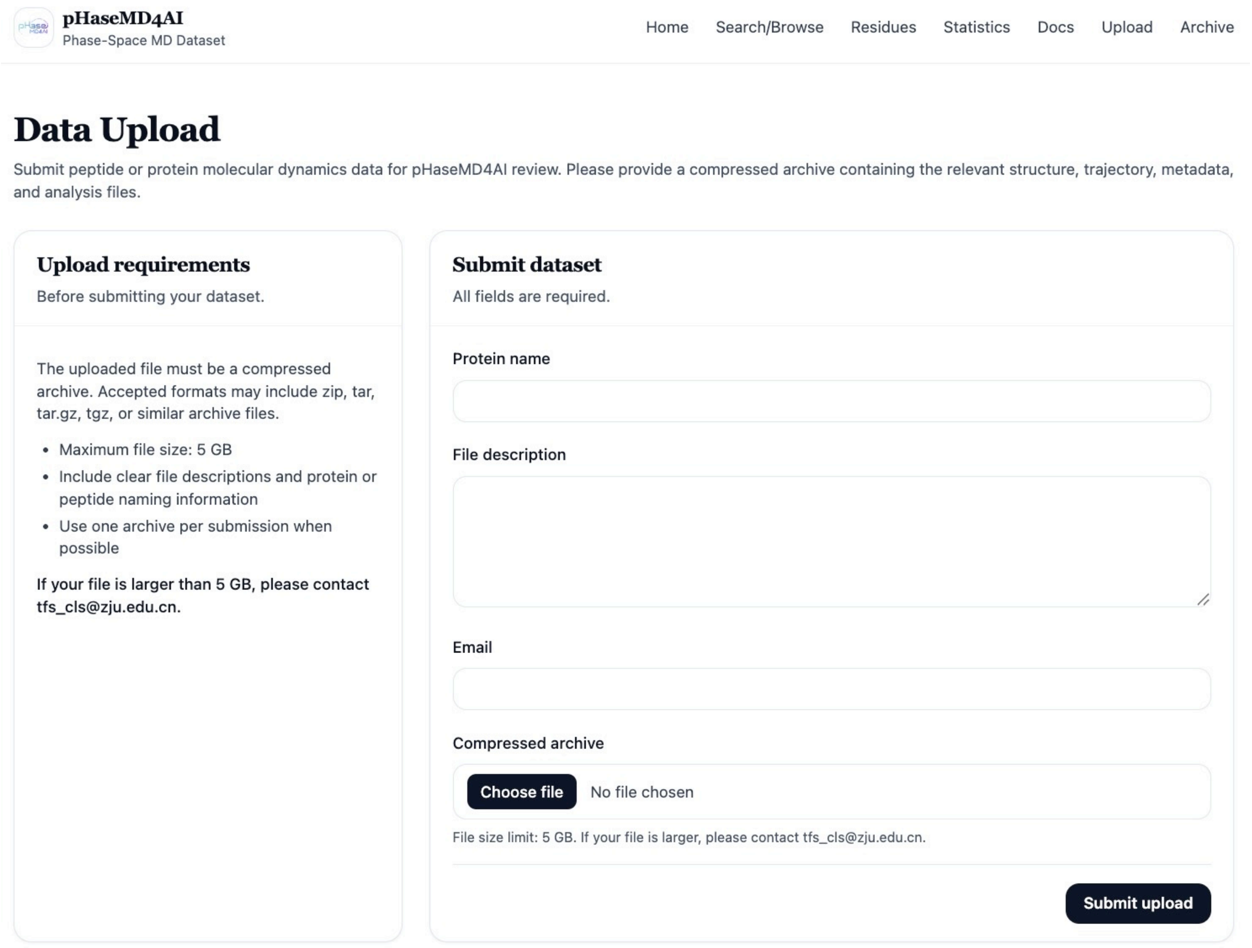


**Figure S20:** Upload page of the pHaseMD4AI web platform.

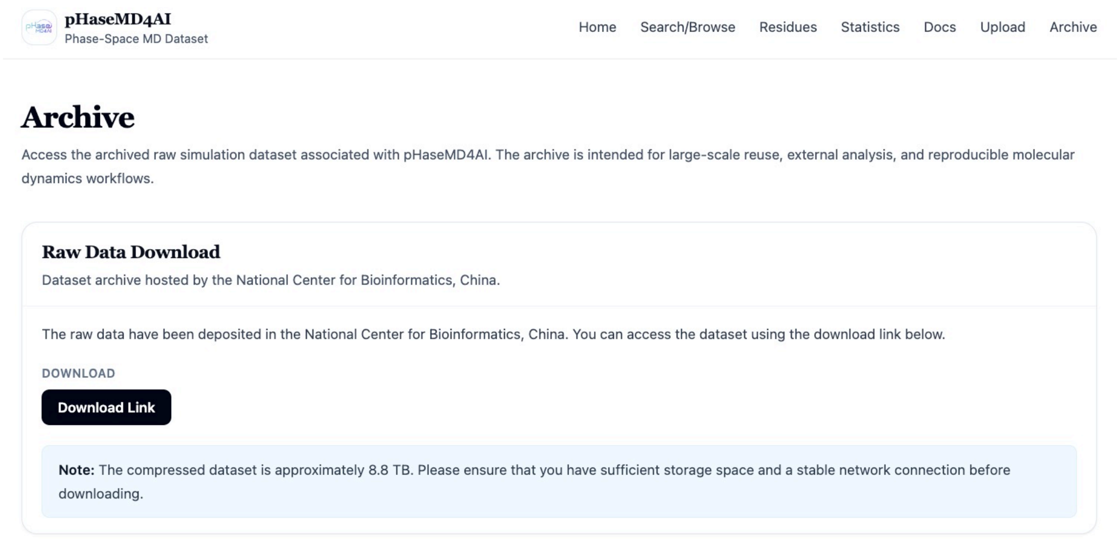


**Figure S21:** Archive page of the pHaseMD4AI web platform.

Zarandi, M.; Sercu, T.; Candido, S.; others Evolutionary-scale prediction of atomic-level protein structure with a language model. *Science* **2023**, *379*, 1123–1130.

(3) Lin, J. Divergence measures based on the Shannon entropy. *IEEE Transactions on Information Theory* **1991**, *37*, 145–151.
